# *N*-Glycosylation of MRS2 balances aerobic and anaerobic energy production by reducing rapid mitochondrial Mg^2+^ influx in conditions of high glucose or impaired respiratory chain function

**DOI:** 10.1101/2024.07.09.602756

**Authors:** Min Peng, Neal D. Mathew, Vernon E. Anderson, Marni J. Falk, Eiko Nakamaru-Ogiso

## Abstract

*N-*linked glycoproteins function in numerous biological processes, modulating enzyme activities as well as protein folding, stability, oligomerization, and trafficking. While *N-*glycosylation of mitochondrial proteins has been detected by untargeted MS-analyses, the physiological existence and roles of mitochondrial protein *N-*linked glycosylation remain under debate. Here, we report that MRS2, a mitochondrial inner membrane protein that functions as the high flux magnesium transporter, is *N-*glycosylated to various extents depending on cellular bioenergetic status. Both *N*-glycosylated and unglycosylated isoforms were consistently detected in mitochondria isolated from mouse liver, rat and mouse liver fibroblast cells (BRL 3A and AFT024, respectively) as well as human skin fibroblast cells. Immunoblotting of MRS2 showed it was bound to, and required stringent elution conditions to remove from, lectin affinity columns with covalently bound concanavalin A or *Lens culinaris* agglutinin. Following peptide:*N-*glycosidase F (PNGase F) digestion of the stringently eluted proteins, the higher M_r_ MRS2 bands gel-shifted to lower M_r_ and loss of lectin affinity was seen. BRL 3A cells treated with two different *N-*linked glycosylation inhibitors, tunicamycin or 6-diazo-5-oxo-L-norleucine, resulted in decreased intensity or loss of the higher M_r_ MRS2 isoform. To investigate the possible functional role of MRS2 *N-* glycosylation, we measured rapid Mg^2+^ influx capacity in intact mitochondria isolated from BRL 3A cells in control media or following treatment with tunicamycin or 6-diazo-5-oxo-L-norleucine. Interestingly, rapid Mg^2+^ influx capacity increased in mitochondria isolated from BRL 3A cells treated with either *N-*glycosylation inhibitor. Forcing reliance on mitochondrial respiration by treatment with either galactose media or the glycolytic inhibitor 2-deoxyglucose or by minimizing glucose concentration similarly reduced the *N-*glycosylated isoform of MRS2, with a correlated concomitant increase in rapid Mg^2+^ influx capacity. Conversely, inhibiting mitochondrial energy production in BRL 3A cells with either rotenone or oligomycin resulted in an increased fraction of *N-*glycosylated MRS2, with decreased rapid Mg^2+^ influx capacity. Collectively, these data provide strong evidence that MRS2 *N*-glycosylation is directly involved in the regulation of mitochondrial matrix Mg^2+^, dynamically communicating relative cellular nutrient status and bioenergetic capacity by serving as a physiologic brake on the influx of mitochondrial matrix Mg^2+^ under conditions of glucose excess or mitochondrial bioenergetic impairment.

## INTRODUCTION

Mg^2+^ is the most abundant cellular divalent cation. As it forms Mg-ATP complexes essential for a myriad of energetically demanding biochemical processes, appropriate Mg^2+^ levels need to be maintained in all living cells and cellular compartments. Mg^2+^ is coordinated by many oxyanions in addition to ATP, including nucleic acids, tricarboxylic acid cycle (TCA) metabolites, and (poly)phosphate. Equilibration of these complexes in cells and within organelles both buffer and reduce the concentration of uncomplexed Mg^2+^ in the cytoplasm [Mg^2+^]_i_ and in the matrix of mitochondria, [Mg^2+^]_m_. In mammalian cells, [Mg^2+^]_m_ has been estimated to range between 0.2 and 1.5 mM [1, 2] while the total [Mg^2+^]_total_ may be as high as 100-300 mM due to large concentrations present in complexes [3]. Part of the observed variation in [Mg^2+^]_m_ may result from changes in the metabolic state of the mitochondria. Mitochondrial Mg^2+^ transport processes play an important role in the regulation of diverse mitochondrial enzymes including phospho-pyruvate dehydrogenase phosphatase (PDHP), which dephosphorylates and concomitantly activates the pyruvate dehydrogenase complex (PDHc) [4], NAD^+^-isocitrate dehydrogenase [5], and respiratory chain (RC) complex enzymes [6, 7]. Similarly, Mg^2+^ in the cytoplasm activates numerous glycolytic enzymes [8], including phosphofructokinase [9], glyceraldehyde 3-phosphate dehydrogenase [10], enolase[11–13], and pyruvate kinase [14].

The human Mg^2+^ transporter gene *MRS2* (Mitochondrial RNA Splicing 2), located at chromosome 6 (6p22.1-p22.3), encodes a mitochondrial protein distantly related to the bacterial CorA family of Mg^2+^ transporters [15]. Yeast Mrs2p was initially identified as being required for mitochondrial RNA group II intron splicing in yeast [16], a function now understood to be an indirect effect of its role to provide adequate magnesium levels to stabilize the functional ribozyme structure [17]. MRS2 has been shown to be an integral mitochondrial inner membrane protein, where it mediates the influx of Mg^2+^ into the mitochondrial matrix and is required for mitochondrial RC complex I [18, 19]. Lpe10p in yeast is a second structural and functional homologue of MRS2, where both proteins are required for the mitochondrial Mg^2+^ transport capacity necessary to sustain full growth [20].

MRS2 is a structurally conserved protein (**Supplemental Figure S1**) having two transmembrane domains that form a Mg^2+^ selective, high-conductance channel controlling Mg^2+^ influx into the mitochondrial matrix via an intrinsic negative feedback mechanism [21]. The conserved GMN motif contributes to the selectivity and regulation of these Mg^2+^ channels [22, 23] and a single consensus *N-*glycosylation sequence (N-X-T/S/C) exists in mouse, rat and human MRS2 (**Supplemental Figure S2**). MRS2 expression has been associated with the maintenance of proper steady-state concentrations of [Mg^2+^]_m_ [3]. As proliferating cells require more Mg^2+^ than quiescent cells, a lack of available Mg^2+^ or defects in Mg^2+^ transport systems can affect their proliferation rate [24, 25]. While it is established that MRS2 is a high flux Mg^2+^ channel, its eukaryotic cell role(s) in regulating oxidative phosphorylation (OXPHOS) and in the pathogenesis of primary mitochondrial diseases (PMD) still remains unknown [26] although 15 separate missense mutations are reported in the ClinVar database as being of undetermined significance [27].

Protein *N-g*lycosylation is a common eukaryotic post-translational modification capable of altering a protein’s activity, stability, localization, and interactions [28]. Although the glycosylation of secreted, membrane, and nucleo-cytosolic proteins has been well-studied, and *O-*glycosylation of mitochondrial proteins is well established [29, 30], whether and how the mitochondrial proteome is *N-*glycosylated is largely unknown [31–34]. *N-*linked glycoproteins are characterized by a large complex saccharide comprised of a common initial sequence of two *N-*acetylglucosamine sugars followed by three mannose moieties in glycosidic linkages (GlcNac_2_Man_3_) linked to an asparagine amide. Protein *N*-glycosylation plays important roles in various cellular processes including cell-to-cell recognition, growth, differentiation, and programmed cell death. Specific *N-*linked glycoprotein changes are associated with disease progression, where identification of *N-*linked glycoproteins may aide disease diagnosis, prognosis, and prediction of treatment outcomes [35]. As *N-*linked glycosylation of mitochondrial proteins was previously thought to be absent, little attention has been paid to its characterization and possible functional significance. However, in yeast suppression of *ALG6* and *ALG7,* which encode proteins required for *N-*glycosylation, result in mitochondrial defects [36]. In addition, NGLY1, a human enzyme responsible for *N*-linked deglycosylation, has been shown to be essential for proper mitochondrial function [37–39], suggesting that regulation of *N*-glycosylation of mitochondrial proteins is important for mitochondrial function. The presence of mitochondrial *N-*glycosylation is supported by an unbiased proteomic study identifying 30 mitochondrial proteins from *Saccharomyces cerevisiae* that bound to lectins concanavalin A (ConA) and wheat germ agglutinin. Interestingly, one of the yeast proteins detected by this global analysis was the mitochondrial inner membrane Mg^2+^ transporter, Lpe10p [40]. The mitochondrial localization was further confirmed by immunoblot analysis where *N-* glycosylation was detected by it having increased electrophoretic mobility following treatment with peptide:*N-*glycosidase F (PNGase F), an enzyme that catalyzes the cleavage of *N-*linked oligosaccharides [41]. It has been gradually accepted that protein glycosylation could contribute to mitochondrial protein localization and function: excessive protein glycosylation of mitochondrial F1Fo-ATP-synthase (complex V) was reported to lead to cell damage and secretory alterations in pancreatic β-cells [42]; the mitochondrial glutamine transporter, SLC15A, was shown to be *N*-glycosylated in pancreatic cancer cells [43]; as well as the mitochondrial NAD^+^ transporter in *Leishmania braziliensis* [44].

Intrigued by the fact that Lpe10p, a close yeast homolog of MRS2, was found to be an *N-*glycosylated mitochondrial protein, we initiated this investigation. Here, we report for the first time that MRS2 localized to mammalian mitochondria exists in either *N-*glycosylated and non*-*glycosylated states. We consistently found both isoforms in isolated mitochondria from mouse liver, rat and mouse liver fibroblast cells (BRL 3A and AFT024, respectively), and human skin fibroblasts. We further found that increases in the fraction of *N-*glycosylated MRS2 isoforms reduced the mitochondrial rapid Mg^2+^ influx capacity, and that the fraction of *N-*glycosylated MRS2 correlates with the relative contributions of OXPHOS and glycolysis to meeting cellular energy demand. We also observed that the *N-*glycosylated MRS2 isoform is increased in several mitochondrial RC disease patient fibroblast cell lines (FCLs). To our knowledge, this is the first report implicating a functional role of *N-*linked glycosylation of a mitochondrial protein and demonstrating its association with both cellular nutrient status and mitochondrial respiratory chain disease.

## METHODS

### Isolation and purification of mitochondria from frozen mouse liver tissues

Previously isolated frozen mouse livers were cut into pieces on ice and transferred into a chilled glass-Teflon homogenizer. Three tissue volumes of cold mitochondrial isolation buffer (MIB: mannitol 23.96 g/L, MOPS 1.04 g/L, pH 7.4, 5 mM EDTA, 1% BSA containing protease inhibitors) was added and the tissue homogenized with 15 strokes. Centrifuging at 750×*g* for 5 min yielded a supernatant, which was centrifuged at 10,000×*g* for 10 min. The pellet was washed twice with MIB after centrifuging at 10,000×*g* for 10 min to yield crude mitochondria. The mitochondrial pellet was then resuspended with 3 mL 15% Percoll^®^ in 16 mL ultracentrifuge tubes, after which 7 mL 23% Percoll and 4 mL 40% Percoll were added from the bottom. The purified mitochondria band (at the interface between 23% and 40% Percoll) was collected and washed twice with MIB after centrifugation at 30,000×*g* for 15 min at 4 ℃. The protein concentration of the mitochondrial fraction was measured by using a BCA protein assay kit (Pierce), after which the mitochondria were stored at -80 ℃.

### Mitoplast preparation from purified mitochondria

Recrystallized digitonin was solubilized at 1.2% (w/v) in MIB. Purified or partially purified mitochondria were stirred gently on ice, with 0.12 mg digitonin/mg mitochondrial protein and equilibrated on ice for 15 min. This preparation was washed 3 times with 3 volumes of MIB followed by centrifugation at 10,000×*g* for 10 min to yield a mitoplast pellet.

### Cell culture and mitochondria isolation from cultured cells

BRL 3A rat liver fibroblasts and AFT024 mouse liver fibroblasts were purchased from the American Type Culture Collection (ATCC; Manassas, VA). Human primary FCLs were obtained when available and/or established in the Clinical CytoGenomics Laboratory from skin biopsies performed following informed consent at the Mitochondrial Medicine Frontier Program of The Children’s Hospital of Philadelphia (M.J.F., CHOP IRB#08–6177). BRL 3A, AFT024 and FCLs were cultured in a mixture of Dulbecco’s modified Eagle’s medium (DMEM; Thermo Fisher Scientific, Waltham, MA) containing 10% heat-inactivated fetal bovine serum (FBS), 1 g/L glucose, and supplemented with 1 mM sodium pyruvate and 2 mM L-glutamine. Human embryonic kidney 293 (HEK293) cells were cultured in RPMI-1640 medium (Sigma, St. Louis, MO) with 10% FBS. All cells were cultured without antibiotics at 37 ℃ in the presence of 5% CO_2_. FCLs from both patients with mitochondrial dysfunction disease and controls were cultured in 10% FBS, 5.6 mM glucose DMEM for 72 h, then the medium was replaced with 10% FBS, 25 mM glucose DMEM for 24 h. Cells were collected in cold phosphate buffered saline (PBS) supplemented with 0.25% trypsin*-*0.02% EDTA, washed twice with cold PBS at 500×*g* for 5 min, and suspended in 5 volumes of subcellular fractionation buffer (250 mM sucrose, 2 mM HEPES pH 7.4, 10 mM KCl, 1.5 mM MgCl_2_, 1 mM EDTA, 1 mM EGTA, 1 mM DTT and protein inhibitor cocktail [1:100]), passed through a 26G needle 10 times using a 3 mL syringe, and left on ice for 20 min. The cells were initially centrifuged at 600×*g* for 10 min, after which the supernatant was centrifuged at 10,000×*g* for 10 min with the resultant mitochondrial pellet being washed twice with cold subcellular fractionation buffer at 10,000×*g* for 10 min at 4 ℃. Mitochondria yields were determined by total protein assay following resuspension.

### Western immunoblot analyses

Isolated mitochondria or mitoplasts were lysed with 1× Laemmli buffer (Bio-Rad,) with β-mercaptoethanol, denatured at 95 ℃ for 4 min, sonicated 1 min at 50 kHz, then loaded on 4–15% or 4-20% polyacrylamide Tris-glycine gradient gels (Bio-Rad, Hercules, CA). Following electrophoresis, separated proteins were transferred from the gel to nitrocellulose membranes (Bio-Rad), and the blots were blocked with Odyssey blocking buffer (LI-COR Biosciences, Lincoln, NE) for 1 h at room temperature, then incubated with a primary antibody in reaction buffer (1:1 of Odyssey blocking buffer and TBS-0.05% Tween*-*20 buffer) overnight at 4 ℃. Polyclonal rabbit antibodies against a 50-mer synthetic peptide derived from human MRS2 were purchased from Novus Biologicals (NBP2 34200), Abcam (ab90295, no longer available), Alomone labs (ANT-148) and Origene (TA338429). The blots were washed three times with washing buffer (TBS-0.05% Tween*-*20) on the shaker for 5 min at room temperature and reacted with Odyssey IRDye (LI-COR Biosciences) goat anti-rabbit secondary antibodies at 1:10,000 dilution for 30 min at room temperature. The blots were next washed four times with washing buffer on the shaker for 5 min at room temperature and then scanned directly using an Odyssey Infrared Imaging System (LI-COR Biosciences). The ratios of *N-*glycosylated to unglycosylated MRS2 isoforms are determined as the ratio of high M_r_:low M_r_ bands as presented in Table 1. The M_r_ of observed bands was determined by interpolation from the Chameleon IRDye molecular weight standards (LiCOR Biosciences, 928-40000).

**Table 1.**
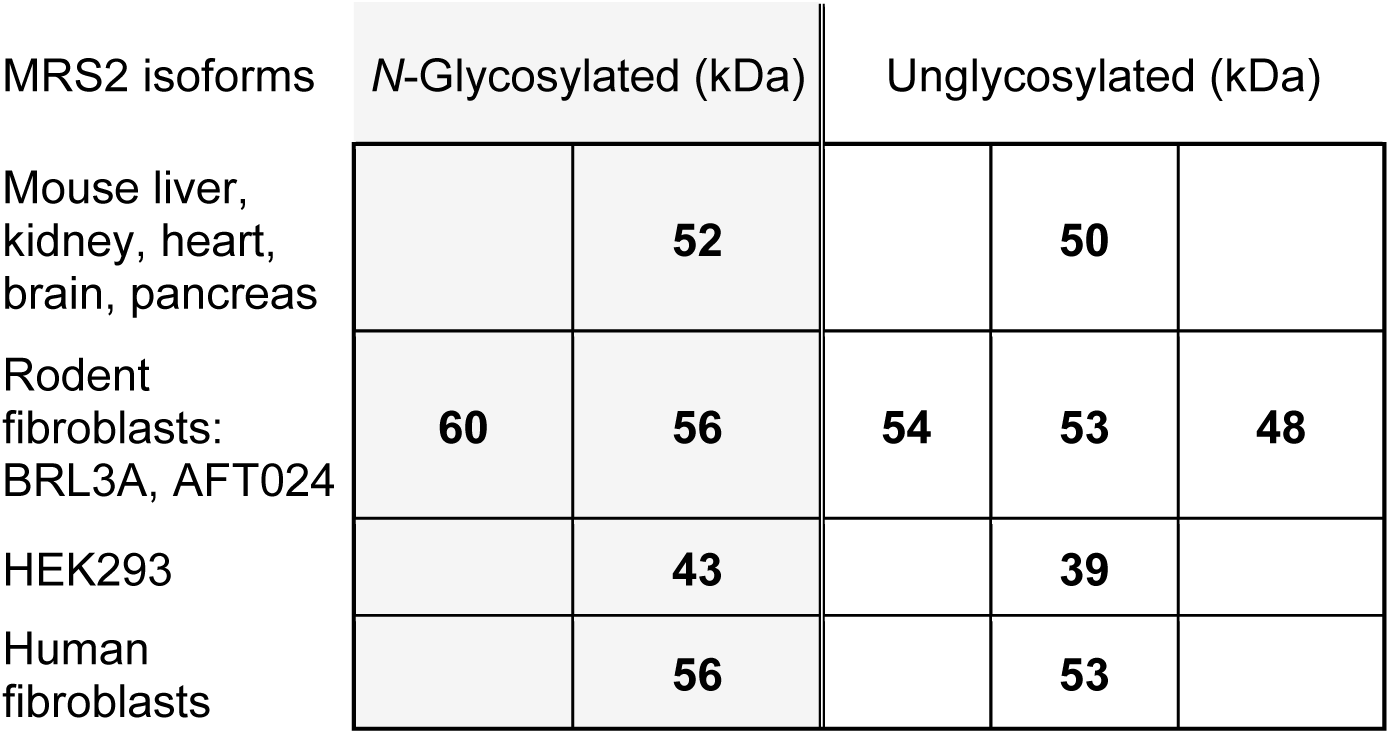

| MRS2 isoforms | N-Glycosylated (kDa) |  | Unglycosylated (kDa) |  |  |
| --- | --- | --- | --- | --- | --- |
| Mouse liver,<br>kidney, heart,<br>brain, pancreas |  | 52 |  | 50 |  |
| Rodent<br>fibroblasts:<br>BRL3A, AFT024 | 60 | 56 | 54 | 53 | 48 |
| HEK293 |  | 43 |  | 39 |  |
| Human<br>fibroblasts |  | 56 |  | 53 |  |

### Lectin*-*glycan binding assay

Lectin (Con A and *Lens culinaris* agglutinin (LCA))-beads were purchased from Vector Laboratories (Burlingame, CA). Initial binding equilibration buffer was 20 mM Tris-HCl, pH 7.4, 1 mM MgCl_2_, 1 mM MnCl_2_, 1 mM CaCl_2_, 150 mM NaCl (0.5 M NaCl for Con A); washing buffer was binding buffer plus 0.1% Tween*-*20; and the stringent elution buffer was 5 mM Tris-HCl, pH 8.0, 150 mM NaCl, 0.05% SDS, 0.5 M D-mannoside (for Con A) or 1% SDS (for LCA). A volume of 200 μL Con A or LCA beads per 1 mg mitochondrial protein, or mitoplasts prepared from 2 mg of mitochondria was used. The lectin*-*beads were washed twice with washing buffer at 500×*g* for 1 min, adding one volume beads of binding buffer and mitochondria in a 1.5 mL microtube and placed on a tube rotator to react at room temperature for 30 min (Con A), or 60 min (LCA). The lectin*-*beads were centrifuged at 200×*g* for 30 s and washed 5 times with washing buffer for 5 min. Lectin*-*binding proteins were recovered with one volume of stringent elution buffer at room temperature or half volume of 2% SDS for 5 min at 95 ℃ on a rotator. Con A-binding glycoproteins recovered with elution buffer were concentrated with Amicon Ultracel-3 kDa filters.

### Expression of *MRS2∷FLAG* in HEK293 cells

*HsMRS2* cDNA was obtained from Genscript (GenScript, NM_020662.3) and the DNA sequence for a FLAG tag fused to the C-terminus [45]. Subsequently Asn96 was conservatively mutated to Gln96 (N96Q) to mutate the lone *N-*glycosylation site in the *MRS2* sequence. HEK293 cells were grown in 100 mm dishes until 90% confluent. A volume of 20 μL Lipofectamine 2000 was mixed with 1 mL Opti-MEM Reduced Serum Medium, and 2 nM *HsMRS2∷FLAG* or human control cDNA, *GRP94* (Origene, NM_003299) was mixed with 1 mL Opti-MEM Reduced Serum Medium, kept at room temperature for 15 min, then Lipofectamine 2000 and lentiviral were added together and were gently mixed.

### Measurement of Mg^2+^ influx capacity of isolated mitochondria

Mitochondria isolated from cultured cells were resuspended with cold [Mg^2+^]_m_ assay buffer (210 mM mannitol, 70 mM sucrose, 2 mM EGTA, 1 mM DTT, 5 mM HEPES-KOH, pH 7.4, 0.5 mM ATP and 1% fraction V BSA) and loaded with 3 μM Mag-Fura-2 AM (Thermo Fisher Scientific, #M1292) at room temperature for 30 min. The mitochondria were washed twice with cold [Mg^2+^]_m_ assay buffer at 1000×*g* for 10 min at 4 ℃. The mitochondrial pellet was suspended with 100 µL [Mg^2+^]_m_ assay buffer. [Mg^2+^]_m_ was characterized by measuring the fluorescence of the probe-loaded mitochondria with a Fluoromax-4 spectrofluorometer (Horiba Scientific, Piscataway, NJ) using two excitation wavelengths, λ_ex-b_ = 340 nm for Mg^2+^-bound and λ_ex-f_ = 380 nm for free Mag-Fura-2 [46] and the single emission wavelength, λ_em_ = 509 nm. Measurements were made before and after the addition of a bolus of exogenous Mg^2+^ as the dissolved chloride salt. All measurements were made at room temperature in 2 mL cuvettes containing 0.5 mg mitochondrial protein in 1 mL of [Mg^2+^]_m_ assay buffer. As adapted from Kolisek at al. [47], after ∼200 s of equilibration time the ratio of fluorescence with 340 nm and 380 nm excitation, [F_exc340/380_]_t=200_, was recorded; exogenous Mg^2+^ was rapidly added to a final concentration of 5 mM and within the mixing time, a large increase in this ratio was observed and recorded [F_exc340/380_]_t=200s_+. The ratio of fluorescence intensities after and before the Mg^2+^ addition, [F_exc340/380_]_t=200s_+**_/_** [F_exc340/380_]_t=200_, we define as the rapid Mg^2+^ influx capacity. In our studies the experimental rapid Mg^2+^ influx was normalized to the control (Eq 1):

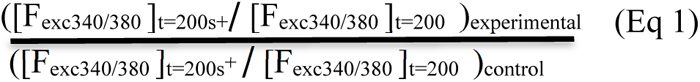

In some assays a final concentration of 1 mM Co(III)Hex (Sigma Aldrich, # 50-180-0331) was added to inhibit Mg^2+^ import but not export, allowing for the slower loss of [Mg^2+^]_m_ and a decrease in F_exc340/380_.

### Knockdown of MRS2 expression in mouse AFT024 cells

AFT024 cells were grown in 100 mm dishes until 90% confluent. A volume of 20 μL Lipofectamine 2000 was mixed with 1 mL Opti-MEM Reduced Serum Medium, and 2 nM *MmMrs2* shRNA), or mouse control shRNA was mixed with 1 mL Opti-MEM Reduced Serum Medium, kept at room temperature for 15 min, then Lipofectamine 2000 and shRNA were added together and were gently mixed. The Lipofectamine 2000-shRNA complex solution (2 mL) was added to the 100 mm dishes containing AFT024 cells in 10 mL of growth medium. A homogenous mixture was obtained by gently rocking the plate. The cells were incubated at 37 ℃ under 5% CO_2_ for 16 h, after which the medium was removed and 2 mL of the Lipofectamine 2000-shRNA complexes added to the incubator for continuous culturing.

## RESULTS

### MRS2 is an *N-*linked glycoprotein

MRS2 is a mitochondrial inner membrane protein that has not been previously recognized to be *N-*glycosylated. However, yeast Lpe10 is a homolog of MRS2 that was found to be *N-*glycosylated by global proteomic analysis [40]. To explicitly examine *N-*glycosylation of mammalian MRS2 by immunoblotting, available antibodies from four commercial sources were investigated to determine their specificity (**Supplemental Figure S3**). Validation of these antibodies was pursued by lentiviral expression in HEK293 cells of FLAG-tagged human MRS2 (right 2 lanes of each panel), where two bands at 43 kDa and 39 kDA were commonly detected by all four antibodies. In the absence of lentiviral overexpression of MRS2, the Abcam and Origene antibodies reproducibly detected these bands. Further, the Novus antibody results corroborate those reported by Joshi and Gohil [48], with minimal native expression detected of the glycosylated 43 kDa band. The presence of MRS2 in isolated mitoplasts (isolated membranes from mitochondria) and purified mitochondria from mouse liver, brain and pancreas consistently appeared in western immunoblots as bands at molecular weights (M_r_) 52 kDa and 50 kDa (**Supplemental Figure 4, Table 1**) while from mouse, rat and human fibroblasts the M_r_ of the *N-* glycosylated an unglycosylated bands appeared at 56 kDa and 53 kDa, respectively (**Figure 1D-E**). The gene structure of human and rodent MRS2 contains between 9 and 12 exons (**Supplemental Figure S1B**), potentially giving rise to different isoforms in different organisms and tissues. In HEK293 cells, two MRS2 isoforms (43 kDa and 39 kDa) were confirmed with lentivirus-MRS2-Flag (**Figures 2A,B, Supplemental Figure 3**), and MRS2 plasmids (data not shown). The use of PNGase-F to quantitatively remove *N-*glycosylation from proteins is well documented [41]. Mitochondrial proteins were extracted from isolated mitochondria and were digested with PNGase F with no digestion as control (to identify *N*-glycosylation), then were analyzed by SDS-PAGE to separate proteins and immunoblotted with anti-MRS2 antibody. A gel shift from 56 kDa to 53 kDa in mitochondrial proteins digested with PNGase F was observed, but there was no gel shift of the 53 kDa MRS2 isoform (**Figure 1D,E**), indicating that only the 56 kDa MRS2 isoform is *N-* glycosylated, and the 53 kDa MRS2 isoform is unglycosylated.

**Figure 1.**
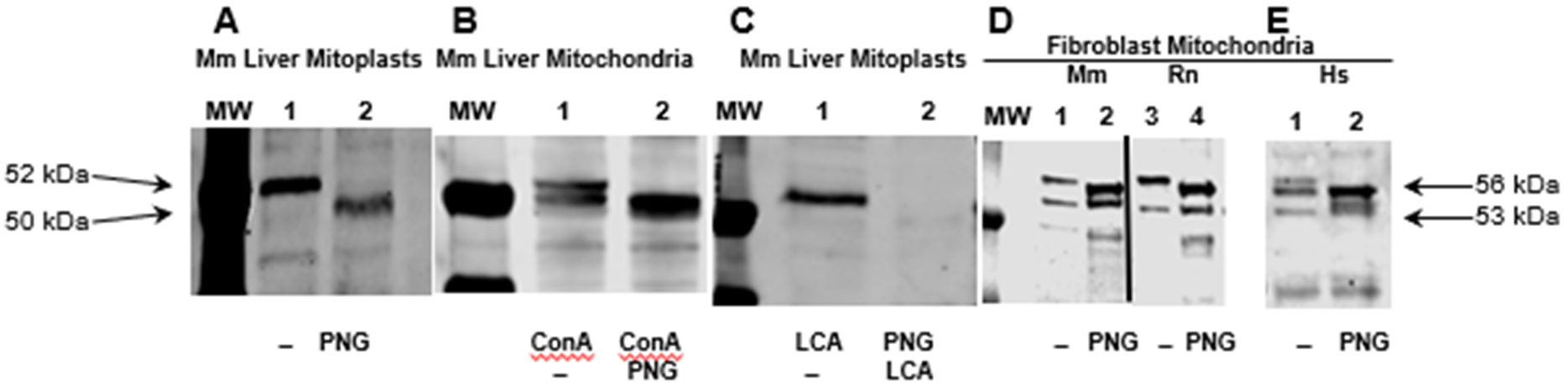
MRS2 is *N*-glycosylated in mouse, rat and human mitochondria. Immunoblot analyses of MRS2 revealed changes following treatment of mitochondrial protein extracts with PNGase F (PNG). **Panels A-E** show 5 separate gels, where unlabeled lanes display molecular weight (MW) markers. **(A)** Proteins isolated from mouse liver (MmL) mitoplasts (MTP) (**1**) before and (**2**) after treatment with PNG. **(B)** Proteins isolated from mouse liver mitochondria (Mito) and affinity purified with conconavalin A (ConA) beads (**1**) before and (**2**) after treatment with PNG. **(C)** Proteins isolated from mouse liver MTP (**1**) without PNG treatment and (**2**) with PNG treatment and then subsequently affinity purified with *Lens culinaris* agglutinin (LCA) beads. **(D)** Proteins isolated from mouse liver AFT024 fibroblast mitochondria (**1**) before and (**2**) after treatment with PNG and proteins isolated from rat (Rn) liver BRL 3A fibroblast mitochondria (**3**) before and (**4**) after treatment with PNG. The vertical solid black line in panel D indicates that the two lanes to its right were not immediately adjacent to the molecular weight control lane (MW). **(E)** Mitochondrial proteins from human fibroblast mitochondria (**1**) before and (**2**) after treatment with PNG. Overall, 52 and 50 kDa bands were detected in Panels A-C, while 56 and 53 kDa bands were detected in panels D and E.

**Figure 2.**
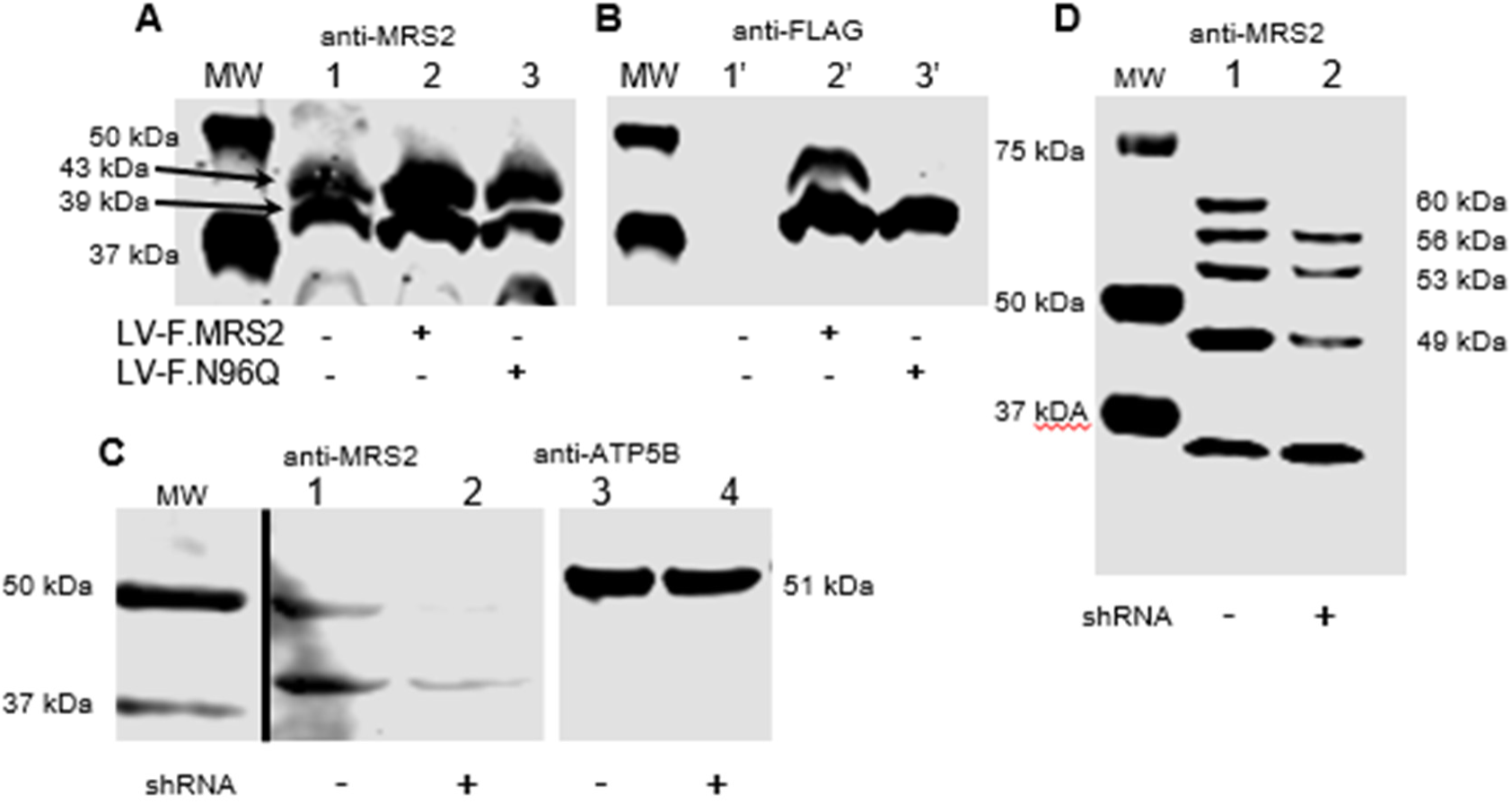
Lentiviral expression of MRS2-FLAG (LV-F.MRS2) in HEK293 cells confirms MRS2 assignment and glycosylation status. Proteins isolated from HEK-293 cells (A) probed with anti-MRS2 and (B) anti-FLAG. (A1) mock-infected cells identify presence of nuclear encoded MRS2 present as two intense bands at 43 and 39 kDa that are (B1’) undetected in the anti-FLAG western blot (A2) expression of wildtype lentiviral MRS2 with anti-MRS2 identifies the same bands and the presence of the same two plasmid-encoded bands are revealed by (B2’) anti-FLAG western blot. (B3) Proteins from HEK-293 cells infected with N96Q MRS2-FLAG reveal the same pattern but in (B3’) the absence of a glycosylation motif resulting from the N96Q mutation causes the higher molecular weight plasmid-encoded band to be undetected. (C,D) A shRNA suppressing MRS2 expression was incorporated into (C) HEK 293 cells and (D) mouse AFT024 cells. The suppression (lanes C2 and D2) relative to control plasmid (lanes C1 and D1) resulted in reproducible reduction of bands associated with both glycosylated and nonglycosylated forms of MRS2. In panel C response to anti-ATP5B was used as a loading control. The AFT024 cells VDAC was used as the loading control (not shown). The vertical solid black lines indicate that the bands shown were not adjacent to the molecular weight standards in the full gel.

We employed two carbohydrate-binding proteins (lectins) with different glycan specificities, Con A, and LCA, to extract glycosylated proteins from purified mouse liver mitochondria. Con A binds specifically to α-D-mannosyl and α-D-glucosyl residues of glycoproteins and peptides [49] while LCA recognizes sequences containing α-linked mannosyl residues and an α-linked fucosyl residue attached to the *N-*acetylchitobiose portion of the core oligosaccharide [50]. The results showed that MRS2 bound to each of the two lectin affinity columns, Con A (**Figure 1B**) and LCA (**Figure 1C**). These lectin binding profiles therefore suggest that MRS2 is a matrix or inner mitochondrial membrane glycoprotein containing mannose and *N-*acetylglucosamine residues. An additional verification of the efficacy of PNGase F treatment is shown by pretreatment of the mitochondrial proteins prior to LCA affinity purification (**Figure 1, lane C2**). **Table 1** identifies different M_r_ species detected in this work in different species and cell types both pre and post PNGase F treatment.

Additional confirmation of the identity of the bands was sought by both lentiviral overexpression of a FLAG-tagged MRS2 in HEK293 cells and RNAi knockdown of the expression of MRS2 in HEK 293 and AFT024 cells. *N-*linked glycans are formed in eucaryotes by the enzyme-catalyzed transfer of the glycan from a dolichol-linked donor to the asparagine side chain amide N in the canonical sequence *N-*X-(S/T) [51, 52]. Within the MRS2 protein sequence the only occurrence of this sequence is at residues 96-98 (**Supplemental Figure S2A**). A mutant protein containing the conservative Asn_96_ to Gln_96_ (N96Q) mutation would then be predicted to be stable and functional but incapable of *N-* glycosylation. The N96Q and unmutated WT proteins were expressed by lentiviral infection of HEK293 cells and identified by incorporating a C-terminal FLAG-tag [45]. **Figure 2A** shows that the cells that expressed wild-type MRS2 while **Figure 2B** where the gel was probed with anti-FLAG antibodies revealed that the WT appeared as two bands corresponding to the *N-*glycosylated and unglycosylated forms at 43 kDa and 39 kDa respectively, while the N96Q mutant only appeared as the nonglycosylated 39 kDa form. These data strengthen the assignment of the bands as MRS2 and the *N-*glycosylation of the higher M_r_ band.

RNAi knock down of MRS2 in HEK293 and mouse AFT024 cells further confirms the MRS2 assignments. In **Figure 2C and 2D** transfection with a shRNA against MRS2 reduces the expression of the bands at 39 kDa in HEK293 cells and at 60, 54, 52 and 48 kDa in AFT024, confirming their assignment as MRS2 isoforms as reported in **Table 1**.

Inhibition of *N-*glycosylation reduces the glycosylation of MRS2 in BRL 3A rat liver fibroblast cells. BRL 3A cells were cultured in DMEM 10% FBS with 5.6 mM glucose for 48 h, then different doses of tunicamycin (TM), a well-characterized inhibitor of *N-*glycosylation [53] were added with no treatment as control. Cells were collected 24 h later, mitochondria were isolated and the MRS2 isoforms were analyzed by immunoblot with anti MRS2 antibody. An additional band was observed after BRL 3A cells were treated with 0.5 µg/mL or 1 µg/mL TM. When BRL 3A were treated with 2.5 µg/mL TM for 4 h, the glycosylated 56 kDa MRS2 was reduced relative to the 52 kDa isoform (**Figure 3A**). BRL 3A cells were additionally treated with 100 µM 6-diazo-5-oxo-L-norleucine (DON), a glutamine analog which inhibits the formation glucosamine and hence of the requisite glycosylated dolichol substrate [54], for 24 h (**Figure 3B**) resulting in a nearly complete loss of the 56 kDa band. These observations are consistent with the 56 kDa band corresponding to an *N-*glycosylated form of MRS2 and the 53 kDa band being an isoform lacking *N-*glycosylation.

**Figure 3.**
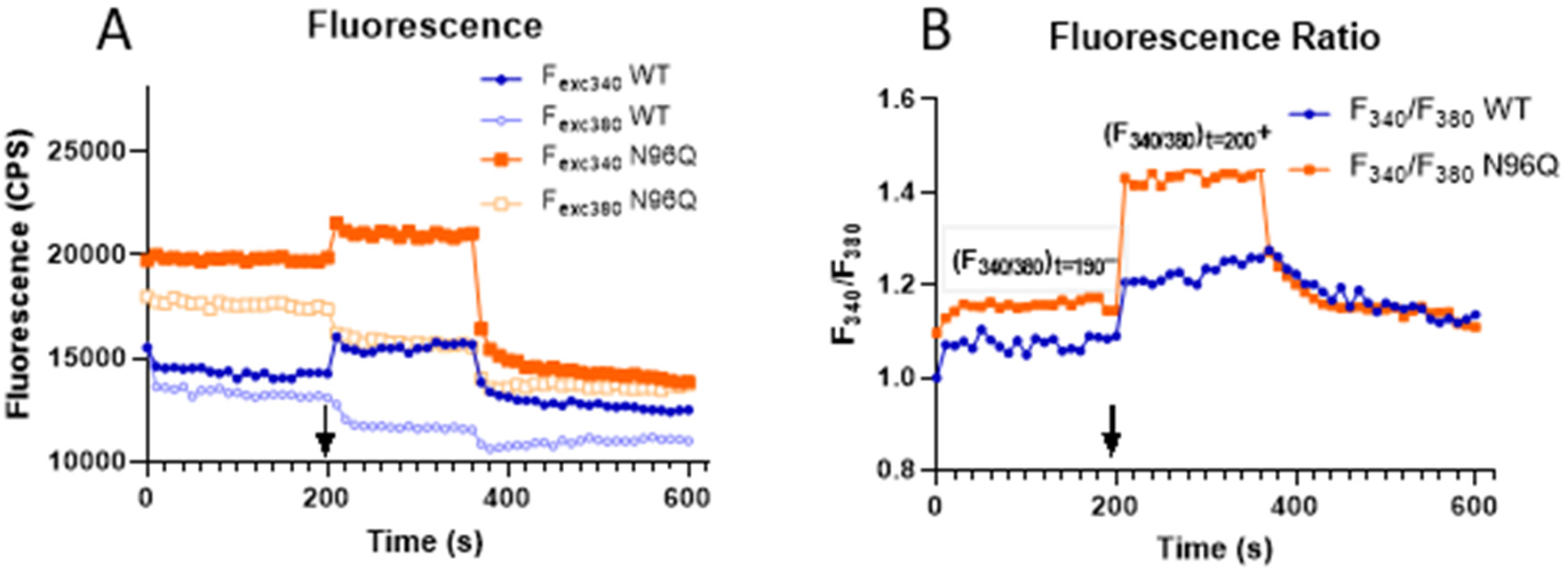
Fluorescence measurement of rapid [Mg^2+^]_m_ influx. (A) Time course of Mag-Fura-2 fluorescence. At 200 s a bolus of 5 mM exogenous Mg^2+^ was added to a suspension of mitochondria (↓) isolated from HEK293 cells following transfection with WT MRS2 (blue) with the MRS2 N96Q mutant (orange) pre-equilibrated with Mag-Fura-2 AM to incorporate the fluorescent Mg^2+^ reporter into the mitochondrial matrix. The increased fluorescence with 340 nm excitation (F_exc340,_ filled symbols) attributed primarily to Mg^2+^ bound form of Mag-Fura-2 contrasts with the decrease in fluorescence with 380 nm excitation (F_exc380_, open symbols) attributed primarily to the free form. (B) The ratio of fluorescence (F_340/380_) amplifies the fluorescence change and reports directly on the [Mg^2+^]_m_ where F_340/380_ is plotted. The F_340/380_ immediately following the addition of 5 mM Mg^2+^ to the initial, (F_340/380_)_t=200_^+^/(F_340/380_)_t=190_-, is a measure of the MRS2-dependent Mg^2+^rapid influx capacity. As it is not possible to assign an absolute concentration, our experiments all report a ratio of experimental to control or in this case mutant to WT: 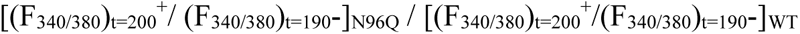 HEK293 cells expressing an *N-*glycosylation deficient N96Q MRS2 mutant displayed a significantly enhanced rapid Mg^2+^ influx capacity. At longer times (∼360 s) an addition of Co(III)hexamine is added to inhibit MRS2 which results in a decrease in the [Mg^2+^]_m_

### Mg^2+^ influx into isolated mitochondria can be measured by spectrofluorometry and is affected by *N-*glycosylation of MRS2

The Mg^2+^ specific fluorescent dye, Mag-Fura-2 reports on free Mg^2+^ concentration by virtue of a shift in the excitation maximum from 380 nm in the Mg^2+^-free form to 340 nm when Mg^2+^ is coordinately bound when measured at the common emission maximum of 500 nm; the Mg^2+^ dissociation constant is ∼ 2.3 mM under physiological conditions [1]. Consequently the F_340_ / F_380_ excitation ratio follows the [Mg^2+^]_m.._ Mag-Fura-2 can be loaded into isolated mitochondria as the membrane permeable ester (Mag-Fura-2 AM) which is trapped when esterases in the matrix hydrolyze the acetoxymethyl esters. Addition of a bolus of external Mg^2+^ to mitochondria isolated from HEK293 cells results in a rapid (< 10 s) influx of Mg^2+^ increasing the [Mg^2+^]_m_ and consequently increasing the F_340_ and decreasing F_380_ (**Figure 3**); recording the ratio F_340_ / F_380_ amplifies the change enhancing the measurement of [Mg^2+^]_m._ This rapid increase in [Mg^2+^]_m_ is attributed to the Mg^2+^ specific, high flux capacity pore formed by an oligomer of MRS2 [47] recently shown by cryoEM to be a pentamer (**Supplemental Figure S2B**) [55] is quantified by Eq. 1. The addition of Co(III)hexamine inhibits MRS2 [47] corroborating that the rapid increase is due to the presence of MRS2 and that the Mag-Fura-2 is reporting on [Mg^2+^]_m_.

MRS2 is the major transport protein for mitochondrial magnesium uptake from cytosol, and the expression of MRS2 is essential for the maintenance of mitochondrial function [3, 18, 20, 47, 56]. After demonstrating that MRS2 was *N-* glycosylated at asparagine-96 the potential role of this glycosylation in Mg^2+^ transport was examined in mitochondria isolated from HEK293 cells expressing the nonglycosylated N96Q mutant. The reduced presence of *N-*glycosylation in this mutant resulted in a large increase in the magnitude of the rapid increase in [Mg^2+^]_m_ (**Figure 3**). This rapid increase in [Mg^2+^]_m_ we identify as the “rapid Mg^2+^ influx capacity”. Thus, the larger rapid Mg^2+^ influx capacity in N96Q glycosylation-deficient mutant implies that the greater *N-*glycosylation present in WT results in a diminished rapid Mg^2+^ influx capacity.

The effects of *N-*glycosylation on MRS2 in BRL 3A cells was further examined by inhibiting the glycosylation with either tunicamycin or DON. The presence of either inhibitor for 24 hours resulted in a decreased relative amount of the *N-* glycosylated 56 kDa band (**Figure 4**). These data further validate the assignment of the 56 kDa band to an *N-*linked glycosylated form of MRS2.

**Figure 4.**
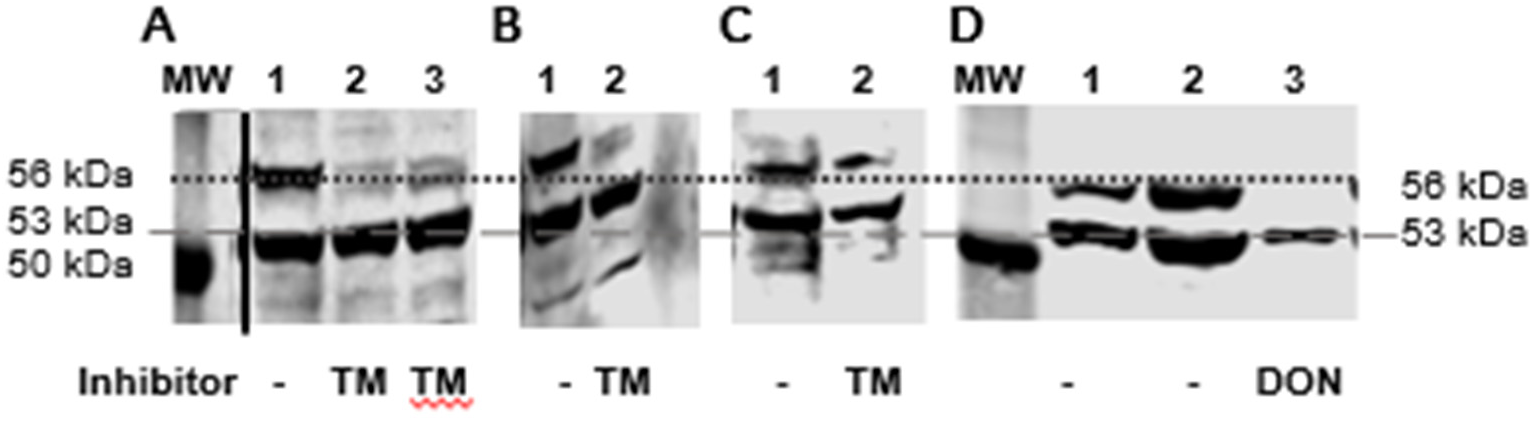
Inhibitors of *N-*glycosylation reduce the ratio of *N*-glycosylated to nonglycosylated MRS2 in BRL 3A cells. The dotted line corresponds to a ∼ MW of 56 kDa assigned to an *N-*glycosylated isoform of MRS2 and the dashed line corresponds to a MW of ∼ 53 kDa and is assigned to a nonglycosylated MRS2 isoform. The MW marker band in the left lanes of Panels A and D is assigned a MW of 50 kDa. (**A, B**) Treatment of BRL 3A cells with tunicamycin (TM) for either (A) 24 h at 0.5 µM (B) 24 h at 1.0 µM or (C) 4 h at 2.5 µM or (D) treatment with DON at 100 µM for 24 h, all with companion controls (-). On each of the four separate gels presented in panels A-D, SDS-PAGE immunoblotting with anti-MRS2 revealed a reduction in the 56 kDa/53 kDa intensity ratio. The efficacy of TM varied with both time and concentration while notably the treatment with DON resulted in a near total loss of the *N-*glycosylated 56 kDa band. The vertical solid black line in panel A indicates that the three lanes shown were not immediately adjacent to the molecular weight marker lane (MW).

### Modulation of glycolysis directly correlates with the *N-*glycosylation status of MRS2

Mammalian ATP generation is largely supported by glycolysis, and OXPHOS. In the following we enhanced glycolysis through increasing glucose availability, and we disfavored either glycolysis or OXPHOS by incubating cells with well-characterized inhibitors of each pathway. In **Figure 5A** and **5B**, BRL 3A cells were cultured in either low glucose (5.6 mM) or high glucose (25 mM) medium for 48, 72 or 96 h; mitochondria were isolated and the mitochondrial MRS2 isoforms were analyzed by immunoblot with anti MRS2. Two important observations from these data points are that high glucose favors *N*-glycosylation and that the differences in *N-*glycosylation become more apparent over the 96 h time span. An alternative method of limiting glucose concentration is to culture the cells at different initial cell densities without replenishing the glucose in the media (**Figure 5C**). At lower cell densities the glucose remains higher throughout the incubation; in the progression from low to high glucose (from an initial 3 M cells to 0.25 M cells) the extent of *N*-glycosylation was observed to increase from barely detectable to ∼50%.

**Figure 5.**
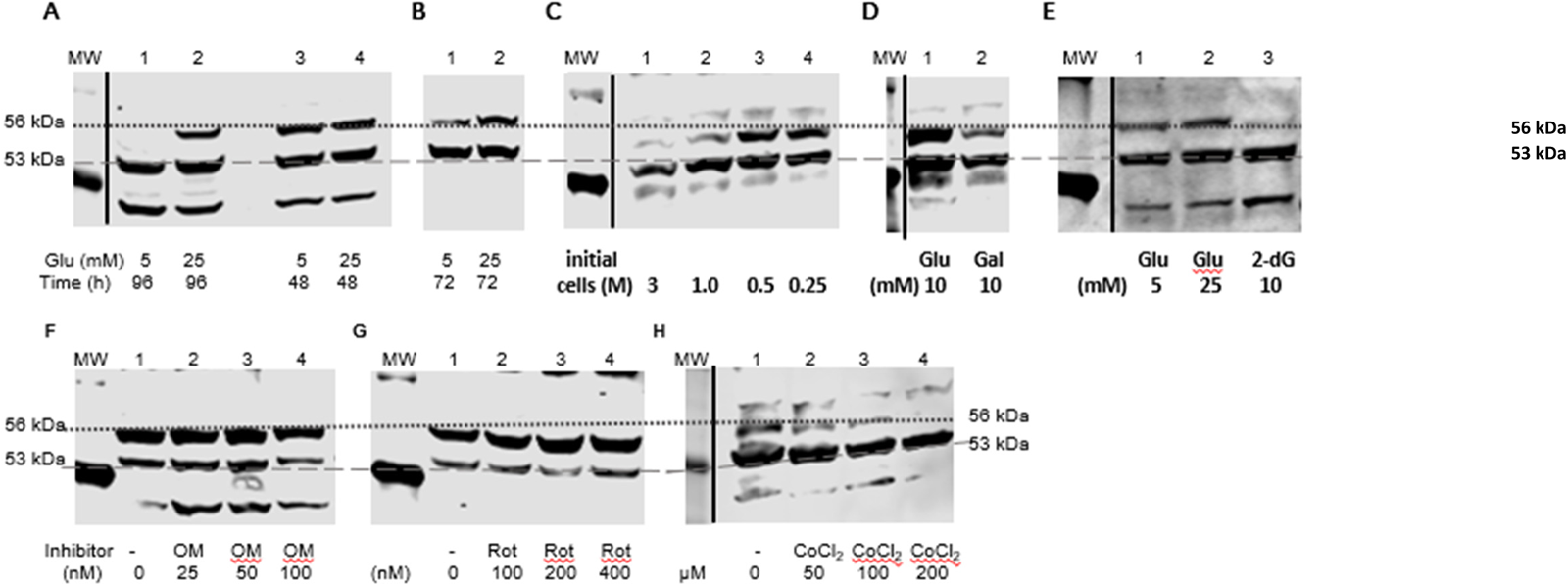
Separately reducing glycolytic or OXPHOS capacity have opposite effects on *N-*glycosylation. Each of the panels represents western immunoblotting results of separate gels with anti-MRS2. The dotted line corresponds to a ∼ MW of 56 kDa and the dashed line corresponds to a MW of ∼ 53 kDa and is assigned to a nonglycosylated MRS2 isoform. The MW marker band in the left lanes is assigned a MW of 50 kDa. (**A,B**) Incubation of (0.5 M initial cell density) BRL3A cells in low (5.6 mM) or high (25 mM) glucose (Glu) for 96, 48 or 72 h. At every time point *N-* glycosylation, and presumably glycolysis, is enhanced at Glu (A2 vs A1, A4 vs A3, B2 vs B1). With time glycosylation is reduced, most apparently at low Glu (A3 to B1 to A1), but also at high Glu (B4 to B2). (**C**) Similarly, the reduction of Glu by altering cell number and hence Glu utilization over 72 h, reduces *N-*glycosylation. Four plates were seeded with 3, 1, 0.5 and 0.25 million (M) cells. The fraction of *N-*glycosylation increased in the same order (lanes C1, C2, C3, C4). Inhibition of glycolysis with either (**D**) galactose (Gal) or (**E**) 2-deoxyglucose (2-dG) similarly reduced the relative amount of *N*-glycosylation, lanes D2 vs D1 and E3 vs E2, respectively. (**F,G**) Incubation of BRL 3A cells with increasing concentrations of oligomycin (OM) (F) or rotenone (Rot) (G) decreased the relative amounts of *N-*glycosylation. (H) Inclusion of CoCl_2_, an inducer of a hypoxic response, resembled the results of inhibiting glycolysis. In Panels A,C,D,E and H the solid vertical lines indicate that the MW marker lane was not immediately adjacent to the lanes shown.

Inhibition of glycolysis or OXPHOS generated opposite results on the observed MRS2 *N*-glycosylation. Inhibition of glycolysis by replacing glucose with the galactose or by 2-deoxyglucose dramatically decreased the extent of *N*-glycosylation (**Figure 5D** and **5E**). This was opposed by the effect of inhibiting OXPHOS. The inclusion of rotenone, a complex I inhibitor, or oligomycin, an ATP synthase inhibitor, resulted in enhanced *N*-glycosylation as shown in **Figure 5F** and **5G**. Cobalt chloride (CoCl_2_) induces a hypoxic response (not hypoxia itself) through stabilizing hypoxia inducible factors 1α and 2α [57]. BRL 3A cells were treated with 0, 50, 100, 200 µM CoCl_2_ for 48 h, mitochondria were isolated, and MRS2 isoforms were analyzed by immunoblot. The *N-*glycosylated 56 kDa MRS2 was diminished in a dose response manner as the CoCl_2_ dose increased; the 56 kDa to 53 kDa ratio of MRS2 sharply decreased until undetectable (**Figure 5H**).

In addition to monitoring the effect of inhibiting either glycolysis or OXPHOS on *N-*glycosylation, the effects of each condition on the rapid Mg^2+^ influx capacity in three biological replicates were measured and shown in **Figure 6** where the effects of a fractional increase or decrease in *N-*glycosylation are correlated with fractional changes in rapid Mg^2+^ influx capacity. As an example, the treatment with the glycosylation inhibitor DON enhanced the rapid Mg^2+^ influx (positive dark gray bar) while inhibiting *N*-glycosylation (negative light gray bar). The data shown in **Figure 6** for the seven different conditions reinforce the correlation of *N-*glycosylation with decreased rapid Mg^2+^ influx capacity and increased glycolysis.

**Figure 6.**
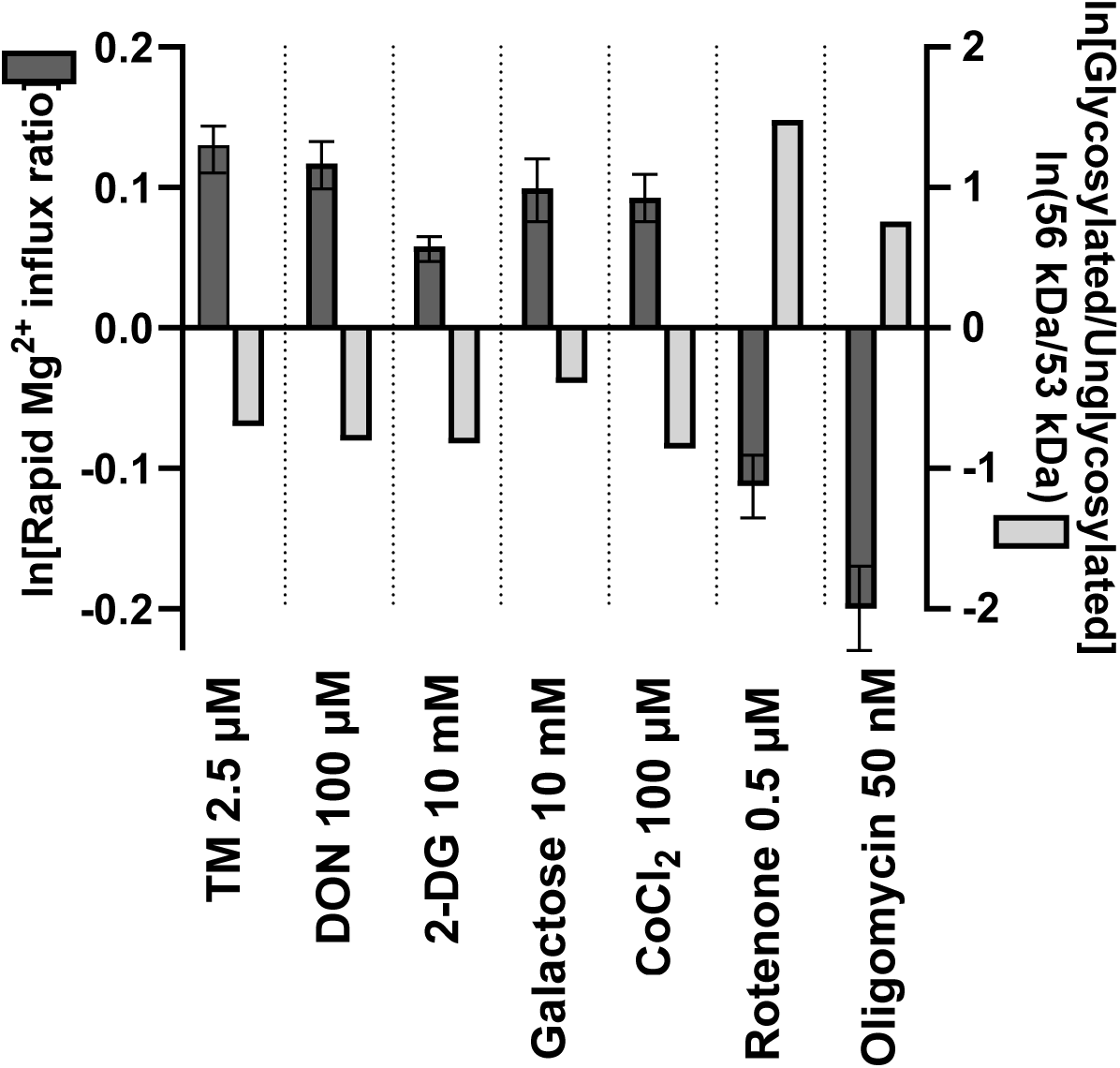
The ratio of the mitochondrial rapid Mg^2+^influx capacity for each of seven experimental conditions (x-axis) normalized by their controls, is displayed as dark gray bars scaled to the left y-axis. (tunicamycin, TM;, DON; 2-deoxyglucose, 2DG) in three biological replicates (mean and standard deviation shown). For these same seven experimental conditions, the ratio of glycosylated to nonglycosylated MRS2 has been determined as the relative intensity of the 56 kDa and 53 kDa bands detected by western immunoblotting (shown in Figures 4-5) and plotted as ln(56 kDa/53 kDa) with the light gray bars scaled by the right y-axis.

### PMD patient FCLs exhibit elevated *N-*glycosylation of MRS2

FCLs were obtained from four patients with different genetically identified PMDs, *C12ORF65* (OMIM #615035, now *MTRFR*) [58], *FBXL4* (OMIM #615471) [59, 60], NDUFS8 (OMIM #618222) [61, 62], and TRIT1 (OMIM #617873) [63]. The results showed the relative protein expressions of 56 kDa to 53 kDa MRS2 isoforms was increased in all four patient cell lines to all four of the control cell lines (**Figure 7**). The rapid Mg^2+^ influx capacity in one patient cell line was reduced compared with the normal cell line consistent with the correlation with small molecule inhibitors (data not shown).

**Figure 7.**
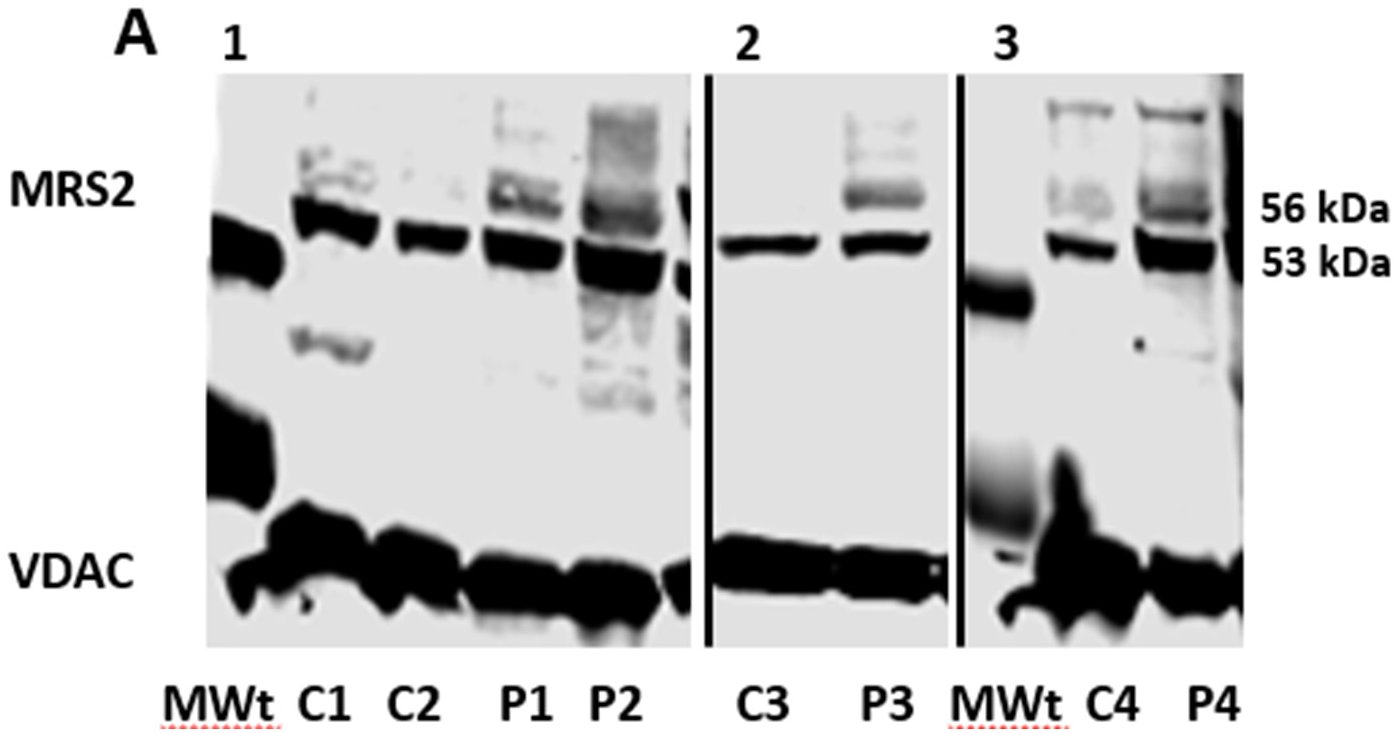
PMD patient fibroblasts show enhanced MRS2 *N*-glycosylation. Incubation of fibroblasts from four separate control (C1-C4) and PMD patients (P1-P4) in 25 mM glucose over 24 h prior to isolating mitochondria and subsequent SDS-PAGE and immunoblotting revealed a consistent pattern of significant MRS2 *N-*glycosylation in the PMD patients’ FCL mitochondria. From three separate PAGE experiments four different PMD patient cell lines were examined. In panel A, FCLs from individuals P1 and P2 with genetically confirmed C12ORF65 and FBXL4 mutations, respectively, were analyzed. In gel B, an FCL from an individual, P3, with genetically confirmed NDUFS8 mutation was analyzed and in gel C an FCL from an individual, P4, with a TRIT1 mutation was analyzed. Different control cell lines, C1-C4 were grown, and mitochondria isolated in parallel. MW standard lane was included at left, the apparent band is 50 kDa. In all four patient cell lines, the *N*-glycosylated form (56 kDa band) is of greater intensity than the nonglycosylated band (53 kDa) when compared to any of the four control cell lines. VDAC was included as a loading control.

## DISCUSSION

Here we have shown varied isoforms of MRS2 are present in rodent tissues and fibroblasts, and that this observation extends to human fibroblasts. In analogy to the N-glycosylation of the homologous yeast Lpe10p, we investigated whether any of the observed isoforms of MRS2 could be attributed to *N-*glycosylation. The selective retention of MRS2 isoforms by lectin affinity columns, the induction of gel-shifts by treatment with PNGase F, the reduction in gel shifted isoforms by treatment of fibroblasts with N-glycosylation inhibitors all support the presence of *N-*glycosylated isoforms of mammalian MRS2. As a single consensus sequence for N-glycosylation is present in the human and rodent MRS2 sequence, glycosylation of this site was tested by lentiviral expression of the FLAG-tagged N96Q mutant. The absence of glycosylation of the mutant (Figure 2B lane 3’) supports the N-glycosylation of the mitochondrial matrix-facing domain of MRS2. To determine if N-glycosylation affects its function, the rapid Mg^2+^ uptake capacity was determined to be significantly increased in isolated mutant mitochondria (Figure 3). Further experiments examined the role of bioenergetics in affecting the in vivo N-glycosylation status, revealing that glucose availability was correlated with N-glycosylation while mitochondrial oxidative phosphorylation activity was inversely correlated (Figure 6). These key results inform our working model of the functional role of MRS2 N-glycosylation in Figure 8.

**Figure 8.**
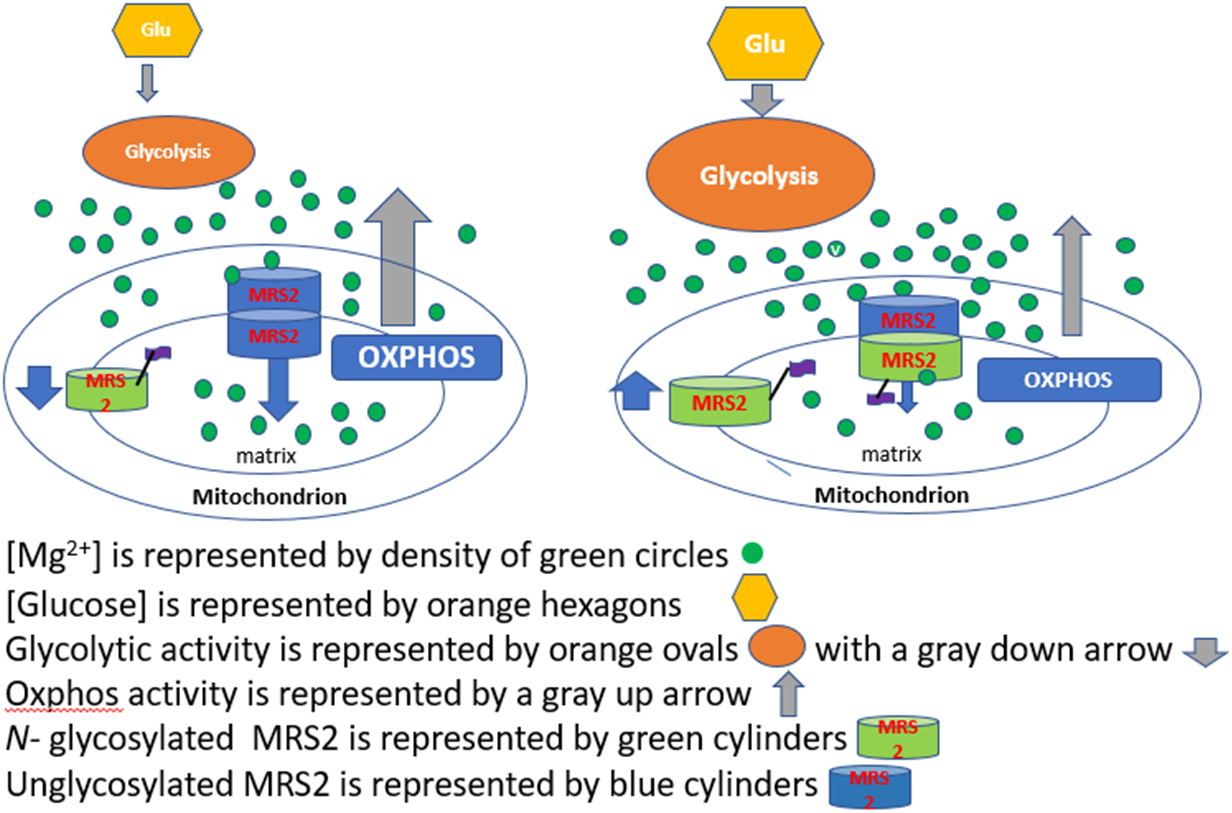
Functional role of *N-*glycosylation of MRS2. The N-glycosylation of a population of MRS2 monomers reduces the set point for rapid Mg^2+^ transport and results in a decrease in the [Mg^2+^]_m_. The decrease in [Mg^2+^]_m_ reduces OXPHOS and secondary effects would enhance glycolysis. Our data show that these effects are reciprocal: enhancing glycolysis increases MRS2 *N-*glycosylation while inhibition of glycolysis decreases MRS2 *N-*glycosylation. PMD patient fibroblasts showed that a decrease in OXPHOS results in enhanced MRS2 *N-*glycosylation in these cells, which would enhance glycolysis.

As a high-capacity ion transporter, a small number of pores per mitochondrial would suffice to maintain the steady state [Mg^2+^]_m_ resulting in a low expression rate [15]. Consequently, MRS2 has been rarely detected by unbiased proteomic methods, with the most certain detection coming from the mitochondrial specific APEX-methodology [64] as noted in Mitocarta3.0 [65]. Our approach, following that of others [41–44] is to either enrich or deplete protein isolates of *N-* glycosylated proteins by affinity purification with lectins or treatment with PNGase F, respectively, and to detect the differences by western immunoblotting. The polyclonal antibodies generated against a human MRS2-derived synthetic peptide used in these procedures lacks specificity as shown in the full gels of four of the five commercially available polyclonal antibodies against MRS2 in **Supplemental Figure S3**. The human MRS2 cDNA sequence data show there are three potential isoforms of human MRS2 with M_r_ between 46.5 kDa and 42 kDa when the proteolyzed mitochondrial leader sequence is omitted [66]. Our data show there are 53 kDa and 56 kDa species by polyacrylamide gel analysis of MRS2 in rat, mouse and human fibroblasts, but 39 kDa and 43 kDa species of MRS2 in HEK293 cells. The binding to the lectin affinity matrices and subsequent gel shift to lower M_r_ following PNGase F treatment provides strong evidence that the 56 kDa isoforms are *N-*glycosylated rodent and human fibroblasts (**Figure 1**). In HEK293 cells the identity of the 39 kDa and 43 kDa bands as authentic MRS2 is further confirmed by the expression and detection of the C*-*terminal FLAG tagged MRS2 at the same M_r_ (**Figure 2A**). The absence of the higher M_r_ band in the N96Q FLAG-tagged mutant in the α-FLAG immunoblot (**Figure 2B**) is strong evidence for MRS2 being solely *N-*glycosylated at Asn_96_. The results from inhibiting MRS2 expression by transfection of mouse fibroblasts with a shRNA corroborated the assignment of the fibroblast proteins detected at 56 and 53 kDa as being MRS2 isoforms **Figure 2D**. The presence of higher M_r_ bands (> 80 kDa) is suggestive of unresolved oligomers, potentially explained by adventitious crosslinking of MRS2 when present as a pentamer in the mitochondrial inner membrane in mammalian cells [55].

### Physiological bioenergetics affect the relative *N-*glycosylation of MRS2

Due to similar intra- and extra-cellular concentrations of Mg^2+^ and their relative constancy, Mg^2+^ variations have been considered based on a presumption of maintaining homeostasis [67] however [Mg^2+^]_m_ has been recently implicated in contributing to the regulation of cellular energy metabolism [56, 68]. Our working model, summarized in **Figure 8** highlight the interconnections between glycolysis, OXPHOS and the regulation of [Mg^2+^]_m_ by MRS2. The consensus view of MRS2 and other Mg^2+^ transporters in the CorA family is that they are effectively chemostatted by [Mg^2+^]_m_, where electrophysiology studies have shown that [Mg^2+^]_m_ bound at the interior allosteric site reduces the probability of these channels being open [23, 69]. Our model shows both the open and closed form of the oligomeric MRS2 pore noting that the recent MRS2 cryo-EM structures are interpreted as depicting the open conformer . The key contribution of our studies is that higher [Mg^2+^]_m_, activation of glycolysis, inhibition of OXPHOS, and *N-*glycosylation of MRS2 are associated with the closed form as our studies indicated that the capacity for glycolysis and OXPHOS to generate ATP inversely affect the set point of the chemostat as the rapid [Mg^2+^]_m_ influx capacity increases when glycolysis is inhibited and decreases when OXPHOS is inhibited. Similarly, the *N-*glycosylation pattern of MRS2 is oppositely affected with *N-*glycosylation increasing when OXPHOS is inhibited, either pharmaceutically or genetically in PMD patients, while inhibition of glycolysis, either pharmaceutically or by its removal from the media, reduces *N-*glycosylation.

Several extensions of this model are tempting to postulate, but not directly addressed by our data. While our model has depicted the *N-*glycosylated form being incorporated into the oligomeric pore, none of our data directly identify the presence of the *N-*glycosylated MRS2 in the oligomer or in any specific oligomeric form. An almost essential role of MRS2 is to prevent the increase in [Mg^2+^]_m_ with increasing membrane potential. With sufficient Mg^2+^ in the cytoplasm, a pore that maintained chemical equilibrium across the inner membrane would result in much higher [Mg^2+^]_m_ while reducing the membrane potential. Thus, it will be interesting to monitor the rapid [Mg^2+^]_m_ influx as a function of membrane potential. Finally, conservation of total Mg^2+^ in the cell would require a decrease in [Mg^2+^]_i_, however the scale of this decrease is difficult to predict because the relative volumes and Mg^2+^ buffering capacities of both compartments are not known.

### Time scale of MRS2 regulation

One aspect of our results that is relevant to the physiological role of MRS2 N-glycosylation is the time required to effect changes. Our experiments were followed for up to 72 h after the initiation of treatment resulting in perceptible changes over time. The long-time frame of regulation is not surprising as most hypothesized pathways for increasing the fraction of *N-*glycosylated MRS2 require either de novo synthesis and glycosylation of MRS2 or the proteolysis of the unglycosylated form, consistent with the half-life of MRS2 in cell culture of human β-lymphocytes determined to be ∼40 h [48]. This contrasts with the rapid [Mg^2+^]_m_ influx in response to cytosolic L-lactate’s induction of release of Mg^2+^ from the endoplasmic reticulum [56]. We suggest that the *N-*glycosylation of MRS2 functions as an adaptation to the cellular environment where enhanced glycolytic capacity (or demand) results in enhanced *N-*glycosylation. This may be a reflection of different roles in different tissues as well as a potential long-term adaptation changes in the cellular environment.

The structures, biogenesis and dynamics of the glycans present on mitochondrial proteins are largely unknown. Similarly it is unknown whether glycosyltransferases and /or glycosidases function in mitochondria and potential roles of glycomic modifications in mitochondrial protein folding, degradation, or function have not been studied. Dolichol phosphate has been found in inner and outer mitochondrial membranes along with the catalytic ability to incorporate sugar residues from their CDP-linked forms. This ability along with the identification of an *N-*glycosyltransferase in both the inner and outer membranes of mouse liver mitochondria [70] suggests that mitochondria may have the ability to synthesize *N-*linked glycoproteins. However to our knowledge direct modification of the matrix side of inner membrane proteins has not been previously identified.

We note that there is only a single potential deglycosylation of MRS2 in the inner mitochondrial membrane if there is an *N-*deglycosylase in the mitochondrial matrix, however to our knowledge and that of others [71] none have been identified. A role for regulation by *N-*glycosylation has been implicated by the identification of a mitochondrial dysfunction present in patients with mutations in the *NGLY1* gene [37–39], a glycosidase that removes *N-*linked glycans from proteins. The mechanism by which deleterious *NGLY1* mutations result in mitochondrial dysfunction are not well characterized, but the most direct cause would be the failure to deglycosylate mitochondrial proteins *in situ* resulting in an altered function. In this work four *NGLY1* deficient fibroblast cell lines demonstrated enhanced *N-*glycosylation of MRS2. However, we have not demonstrated *N-*glycosylated MRS2 is a substrate for NGLY1; to the contrary we have shown that enhanced dependence on glycolysis could similarly enhance MRS2 *N-*glycosylation as a secondary effect.

A common feature of cancer bioenergetics known as the Warburg effect [72], is the reliance on enhanced glycolysis for ATP production. While enhanced glycolysis in certain cancers is due to an impairment of mitochondrial function [73, 74] an important aspect of the Warburg effect is that the function of OXPHOS [75] remains essential. As we have correlated the extent of *N-*linked glycosylation of MRS2 with the capacity of glycolysis and OXPHOS to provide ATP it would be important to investigate whether MRS2 expression and its *N-*glycosylation are involved in cancer mechanisms and potential therapy.

## Contributors

Conceptualization: M.J.F. and E.O.; Funding acquisition: M.J.F.; Protein analysis N.M.; Methodology: M.P. and E.O.; Original draft: M.P., E.O., V.A., M.J.F. All authors reviewed and approved the final manuscript.

## Acknowledgements

We are grateful to Yi Cheng, MD, for her laboratory experimental assistance.

## Ethics Statement

All human subjects research was performed per Children’s Hospital of Philadelphia (CHOP) Institutional Review Board approved study #08-6177 (Falk, PI).

## REFERENCES

1. Rutter, G.A., et al., Measurement of matrix free Mg2+ concentration in rat heart mitochondria by using entrapped fluorescent probes. Biochem J, 1990. 271(3): p. 627–34.

2. Jung, D.W., L. Apel, and G.P. Brierley, Matrix free Mg2+ changes with metabolic state in isolated heart mitochondria. Biochemistry, 1990. 29(17): p. 4121–8.

3. Merolle, L., et al., Overexpression of the mitochondrial Mg channel MRS2 increases total cellular Mg concentration and influences sensitivity to apoptosis. Metallomics, 2018. 10(7): p. 917–928.

4. Thomas, A.P., T.A. Diggle, and R.M. Denton, Sensitivity of pyruvate dehydrogenase phosphate phosphatase to magnesium ions. Similar effects of spermine and insulin. Biochem J, 1986. 238(1): p. 83–91.

5. Qi, F., X. Chen, and D.A. Beard, Detailed kinetics and regulation of mammalian NAD-linked isocitrate dehydrogenase. Biochim Biophys Acta, 2008. 1784(11): p. 1641–51.

6. Yamanaka, R., et al., Mitochondrial Mg(2+) homeostasis decides cellular energy metabolism and vulnerability to stress. Sci Rep, 2016. 6: p. 30027.

7. La Piana, G., et al., Effect of magnesium ions on the activity of the cytosolic NADH/cytochrome c electron transport system. FEBS J, 2008. 275(24): p. 6168–79.

8. Garfinkel, L. and D. Garfinkel, Magnesium regulation of the glycolytic pathway and the enzymes involved. Magnesium, 1985. 4(2-3): p. 60–72.

9. Etiemble, J., et al., Influence of free Mg2+ on the kinetics of human erythrocyte phosphofructokinase. Biochimie, 1981. 63(1): p. 61–5.

10. Molnár, M. and M. Vas, Mg2+ affects the binding of ADP but not ATP to 3-phosphoglycerate kinase. Correlation between equilibrium dialysis binding and enzyme kinetic data. Biochem J, 1993. 293 (Pt 2)(Pt 2): p. 595–9.

11. Anderson, V.E., The mechanism of yeast enolase. 1981: University of Wisconsin--Madison.

12. Farrar, W.W. and W.C. Deal, Jr., Purification and properties of pig liver and muscle enolases. J Protein Chem, 1995. 14(6): p. 487–97.

13. Poyner, R.R., W.W. Cleland, and G.H. Reed, Role of metal ions in catalysis by enolase: an ordered kinetic mechanism for a single substrate enzyme. Biochemistry, 2001. 40(27): p. 8009–17.

14. Nowak, T. and C. Suelter, Pyruvate kinase: activation by and catalytic role of the monovalent and divalent cations. Mol Cell Biochem, 1981. 35(2): p. 65–75.

15. Zsurka, G., J. Gregan, and R.J. Schweyen, The human mitochondrial Mrs2 protein functionally substitutes for its yeast homologue, a candidate magnesium transporter. Genomics, 2001. 72(2): p. 158–68.

16. Wiesenberger, G., M. Waldherr, and R.J. Schweyen, The nuclear gene MRS2 is essential for the excision of group II introns from yeast mitochondrial transcripts in vivo. J Biol Chem, 1992. 267(10): p. 6963–9.

17. Gregan, J., M. Kolisek, and R.J. Schweyen, Mitochondrial Mg(2+) homeostasis is critical for group II intron splicing in vivo. Genes Dev, 2001. 15(17): p. 2229–37.

18. Piskacek, M., et al., Conditional knockdown of hMRS2 results in loss of mitochondrial Mg(2+) uptake and cell death. J Cell Mol Med, 2009. 13(4): p. 693–700.

19. Wolf, F.I., et al., Regulation of magnesium content during proliferation of mammary epithelial cells (HC-11). Front Biosci, 2004. 9: p. 2056–62.

20. Gregan, J., et al., The mitochondrial inner membrane protein Lpe10p, a homologue of Mrs2p, is essential for magnesium homeostasis and group II intron splicing in yeast. Mol Gen Genet, 2001. 264(6): p. 773–81.

21. Schindl, R., et al., Mrs2p forms a high conductance Mg2+ selective channel in mitochondria. Biophys J, 2007. 93(11): p. 3872–83.

22. Sponder, G., et al., The G-M-N motif determines ion selectivity in the yeast magnesium channel Mrs2p. Metallomics, 2013. 5(6): p. 745–52.

23. Dalmas, O., et al., Molecular mechanism of Mg2+-dependent gating in CorA. Nat Commun, 2014. 5: p. 3590.

24. Wolf, F.I. and V. Trapani, Cell (patho)physiology of magnesium. Clin Sci (Lond), 2008. 114(1): p. 27–35.

25. Maguire, M.E. and J.A. Cowan, Magnesium chemistry and biochemistry. Biometals, 2002. 15(3): p. 203–10.

26. Wolf, F.I. and V. Trapani, Multidrug resistance phenotypes and MRS2 mitochondrial magnesium channel: two players from one stemness? Cancer Biol Ther, 2009. 8(7): p. 615–7.

27. MRS2[Gene]. 2024 [cited 2024 03/13/2024]; Clinical Variants of MRS2]. Available from: https://www.ncbi.nlm.nih.gov/clinvar/?term=MRS2[gene].

28. Marth, J.D. and P.K. Grewal, Mammalian glycosylation in immunity. Nat Rev Immunol, 2008. 8(11): p. 874–87.

29. Hu, Y., et al., Increased enzymatic O-GlcNAcylation of mitochondrial proteins impairs mitochondrial function in cardiac myocytes exposed to high glucose. J Biol Chem, 2009. 284(1): p. 547–555.

30. Burnham-Marusich, A.R. and P.M. Berninsone, Multiple proteins with essential mitochondrial functions have glycosylated isoforms. Mitochondrion, 2012. 12(4): p. 423–7.

31. Zhu, J., et al., Comprehensive mapping of protein N-glycosylation in human liver by combining hydrophilic interaction chromatography and hydrazide chemistry. J Proteome Res, 2014. 13(3): p. 1713–21.

32. Stadlmann, J., et al., Comparative glycoproteomics of stem cells identifies new players in ricin toxicity. Nature, 2017. 549(7673): p. 538–542.

33. Han, D., et al., Characterization of the membrane proteome and N-glycoproteome in BV-2 mouse microglia by liquid chromatography-tandem mass spectrometry. BMC Genomics, 2014. 15: p. 95.

34. Fang, P., et al., In-depth mapping of the mouse brain N-glycoproteome reveals widespread N-glycosylation of diverse brain proteins. Oncotarget, 2016. 7(25): p. 38796–38809.

35. Tian, Y. and H. Zhang, Characterization of disease-associated N-linked glycoproteins. Proteomics, 2013. 13(3-4): p. 504–11.

36. Mendelsohn, R.D., et al., A hypomorphic allele of the first N-glycosylation gene, ALG7, causes mitochondrial defects in yeast. Biochim Biophys Acta, 2005. 1723(1-3): p. 33–44.

37. Yang, K., et al., N-glycanase NGLY1 regulates mitochondrial homeostasis and inflammation through NRF1. J Exp Med, 2018. 215(10): p. 2600–2616.

38. Panneman, D.M., et al., Variants in NGLY1 lead to intellectual disability, myoclonus epilepsy, sensorimotor axonal polyneuropathy and mitochondrial dysfunction. Clin Genet, 2020. 97(4): p. 556–566.

39. Kong, J., et al., Mitochondrial function requires NGLY1. Mitochondrion, 2018. 38: p. 6–16.

40. Kung, L.A., et al., Global analysis of the glycoproteome in Saccharomyces cerevisiae reveals new roles for protein glycosylation in eukaryotes. Mol Syst Biol, 2009. 5: p. 308.

41. Tarentino, A.L., C.M. Gómez, and T.H. Plummer, Jr., Deglycosylation of asparagine-linked glycans by peptide:N-glycosidase F. Biochemistry, 1985. 24(17): p. 4665–71.

42. Anello, M., et al., Glucosamine-induced alterations of mitochondrial function in pancreatic beta-cells: possible role of protein glycosylation. Am J Physiol Endocrinol Metab, 2004. 287(4): p. E602–8.

43. Yoo, H.C., et al., A Variant of SLC1A5 Is a Mitochondrial Glutamine Transporter for Metabolic Reprogramming in Cancer Cells. Cell Metab, 2020. 31(2): p. 267–283.e12.

44. Morales Herrera, D.S., et al., Identification and sub-cellular localization of a NAD transporter in Leishmania braziliensis (LbNDT1). Heliyon, 2020. 6(7): p. e04331.

45. Hopp, T.P., et al., A Short Polypeptide Marker Sequence Useful for Recombinant Protein Identification and Purification. Bio/Technology, 1988. 6(10): p. 1204–1210.

46. Froschauer, E.M., et al., Fluorescence measurements of free [Mg2+] by use of mag-fura 2 in Salmonella enterica. FEMS Microbiol Lett, 2004. 237(1): p. 49–55.

47. Kolisek, M., et al., Mrs2p is an essential component of the major electrophoretic Mg2+ influx system in mitochondria. The EMBO Journal, 2003. 22(6): p. 1235–1244.

48. Joshi, A. and V.M. Gohil, Cardiolipin deficiency leads to the destabilization of mitochondrial magnesium channel MRS2 in Barth syndrome. Hum Mol Genet, 2023.

49. Reeke, G.N., Jr., et al., Structure and function of concanavalin A. Adv Exp Med Biol, 1975. 55: p. 13–33.

50. Matsumura, K., et al., Carbohydrate binding specificity of a fucose-specific lectin from Aspergillus oryzae: a novel probe for core fucose. J Biol Chem, 2007. 282(21): p. 15700–8.

51. Welply, J.K., et al., Substrate recognition by oligosaccharyltransferase. Studies on glycosylation of modified Asn-X-Thr/Ser tripeptides. J Biol Chem, 1983. 258(19): p. 11856–63.

52. Hart, G.W., et al., Primary structural requirements for the enzymatic formation of the N-glycosidic bond in glycoproteins. Studies with natural and synthetic peptides. J Biol Chem, 1979. 254(19): p. 9747–53.

53. Elbein, A.D., Inhibitors of the biosynthesis and processing of N-linked oligosaccharides. CRC Crit Rev Biochem, 1984. 16(1): p. 21–49.

54. Telser, A., H.C. Robinson, and A. Dorfman, The biosynthesis of chondroitin-sulfate protein complex. Proc Natl Acad Sci U S A, 1965. 54(3): p. 912–9.

55. Li, M., et al., Molecular basis of Mg(2+) permeation through the human mitochondrial Mrs2 channel. Nat Commun, 2023. 14(1): p. 4713.

56. Daw, C.C., et al., Lactate Elicits ER-Mitochondrial Mg(2+) Dynamics to Integrate Cellular Metabolism. Cell, 2020. 183(2): p. 474–489.e17.

57. Muñoz-Sánchez, J. and M.E. Chánez-Cárdenas, The use of cobalt chloride as a chemical hypoxia model. J Appl Toxicol, 2019. 39(4): p. 556–570.

58. Antonicka, H., et al., Mutations in C12orf65 in patients with encephalomyopathy and a mitochondrial translation defect. Am J Hum Genet, 2010. 87(1): p. 115–22.

59. Gai, X., et al., Mutations in FBXL4, encoding a mitochondrial protein, cause early-onset mitochondrial encephalomyopathy. Am J Hum Genet, 2013. 93(3): p. 482–95.

60. Lavorato, M., et al., Dichloroacetate improves mitochondrial function, physiology, and morphology in FBXL4 disease models. JCI Insight, 2022. 7(16).

61. Guha, S., et al., Pre-clinical evaluation of cysteamine bitartrate as a therapeutic agent for mitochondrial respiratory chain disease. Hum Mol Genet, 2019. 28(11): p. 1837–1852.

62. Guha, S., et al., Combinatorial glucose, nicotinic acid and N-acetylcysteine therapy has synergistic effect in preclinical C. elegans and zebrafish models of mitochondrial complex I disease. Hum Mol Genet, 2021. 30(7): p. 536–551.

63. Yarham, J.W., et al., Defective i6A37 modification of mitochondrial and cytosolic tRNAs results from pathogenic mutations in TRIT1 and its substrate tRNA. PLoS Genet, 2014. 10(6): p. e1004424.

64. Rhee, H.W., et al., Proteomic mapping of mitochondria in living cells via spatially restricted enzymatic tagging. Science, 2013. 339(6125): p. 1328–1331.

65. Rath, S., et al., MitoCarta3.0: an updated mitochondrial proteome now with sub-organelle localization and pathway annotations. Nucleic Acids Res, 2021. 49(D1): p. D1541–d1547.

66. UniProt: the Universal Protein Knowledgebase *in* 2023 Nucleic Acids Research, 2023. **51**(D1): p. D523–D531. https://www.uniprot.org/uniprotkb/Q9HD23/entry#expression.

67. Morrison, A.R., Magnesium Homeostasis: Lessons from Human Genetics. Clin J Am Soc Nephrol, 2023. 18(7): p. 969–78.

68. Madaris, T.R., et al., Limiting Mrs2-dependent mitochondrial Mg(2+) uptake induces metabolic programming in prolonged dietary stress. Cell Rep, 2023. 42(3): p. 112155.

69. Uthayabalan, S., et al., The human MRS2 magnesium-binding domain is a regulatory feedback switch for channel activity. Life Sci Alliance, 2023. 6(4).

70. Levrat, C., et al., Study of the N-glycoprotein biosynthesis through dolichol intermediates in the mitochondrial membranes. Int J Biochem, 1989. 21(3): p. 265–78.

71. *MitoCarta3.0: An Inventory of Mammalian Mitochondrial Proteins and Pathways*. 2020 2020 [cited 2024 March 21, 2024]; Available from: https://www.broadinstitute.org/mitocarta/mitocarta30-inventory-mammalian-mitochondrial-proteins-and-pathways.

72. Koppenol, W.H., P.L. Bounds, and C.V. Dang, Otto Warburg’s contributions to current concepts of cancer metabolism. Nat Rev Cancer, 2011. 11(5): p. 325–37.

73. Chandra, D. and K.K. Singh, Genetic insights into OXPHOS defect and its role in cancer. Biochim Biophys Acta, 2011. 1807(6): p. 620–5.

74. Owens, K.M., et al., Impaired OXPHOS complex III in breast cancer. PLoS One, 2011. 6(8): p. e23846.

75. DeBerardinis, R.J. and N.S. Chandel, We need to talk about the Warburg effect. Nat Metab, 2020. 2(2): p. 127–129.

