## Supplementary Figures for "*N*-Glycosylation of MRS2 balances aerobic and anaerobic energy production by reducing rapid mitochondrial Mg^2+^ influx in conditions of high glucose or impaired respiratory chain function"

### Supplemental Figures

#### Supplemental Figure 1A. MRS2 is highly conserved from human to yeast.

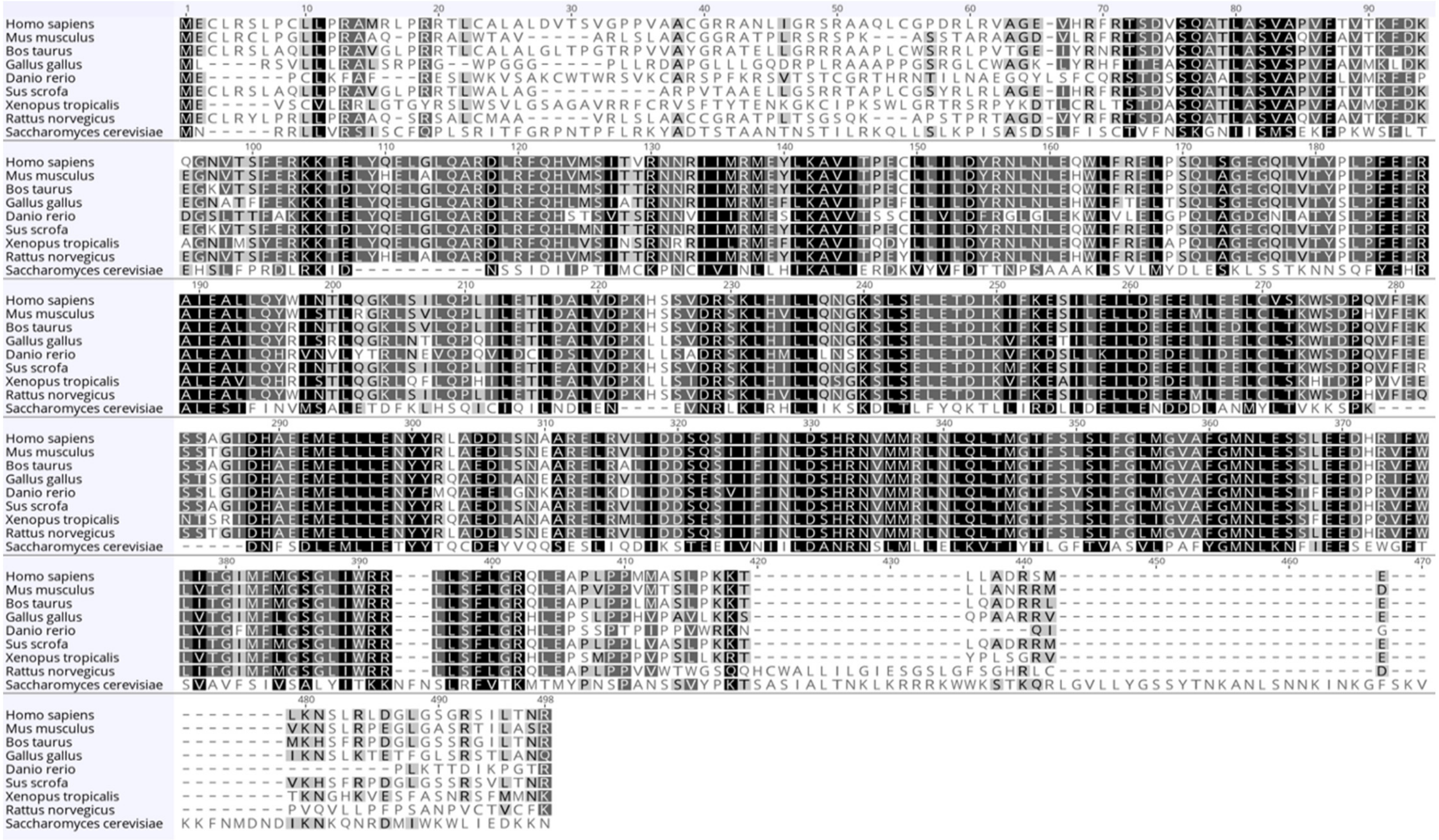


**Supplemental Figure S1A**. Aligned sequences of eight vertebrate MRS2 homologs and yeast Mrs2p. Largely conserved residues are highlighted as white letters with black background, while less well conserved residues have a dark gray background, residues with limited conservation are shown with a light gray background. The blue bar from residue 216 to 265 identifies the antigenic peptide used to generate rabbit polyclonal antibodies, highlighting the conservation of this sequence between human, rat and mouse sequences.

### *Supplemental Figure 1B: human MRS2 gene structure*

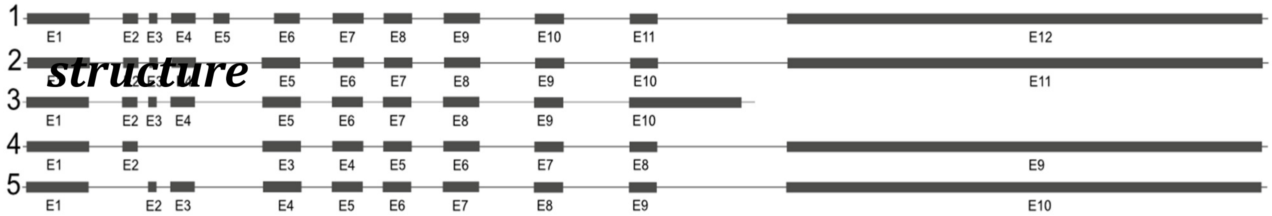


**Supplemental Figure S1B**. Exon structure of human cDNAs mapped onto the chromosome structure. *MRS2* is comprises up to 12 different exons. The existence of five human isoforms complicates the potential identification and pattern of western blots probed for MRS2.

https://www.ncbi.nlm.nih.gov/gene/57380#summary

https://useast.ensembl.org/Homo_sapiens/Gene/Splice?db=core;g=ENSG00000124532;r=6:24402908-24426194

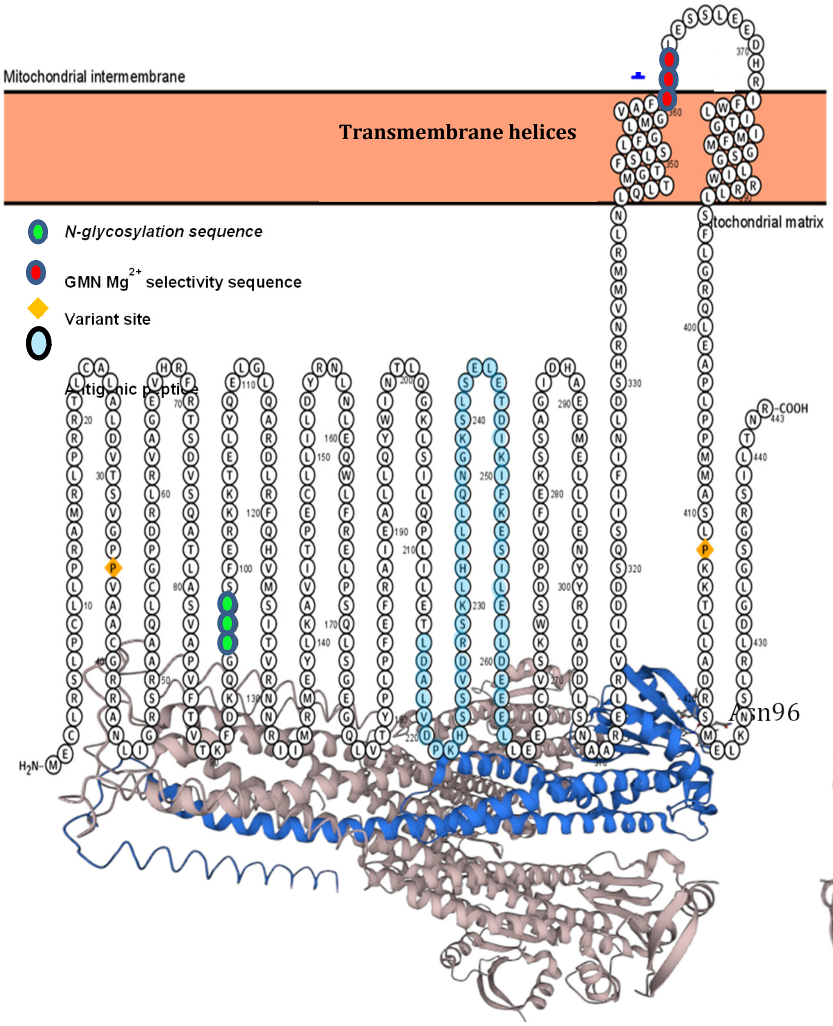


**Sequence organization of the human MRS2 monomer.** The transmembrane orientation of MRS2 is depicted with two transmembrane helices. The regulated pore is a pentamer with intersubunit contacts being made by the helices. The GMN $Mg^{2+}$ selectivity site is identified in red. The single consensus *N*-glycosylation site is highlighted in lime green, two proline residues that were identified as variants are identified as yellow diamonds and the commercial anti-MRS2 antibody was generated with the oligopeptide highlighted in blue.

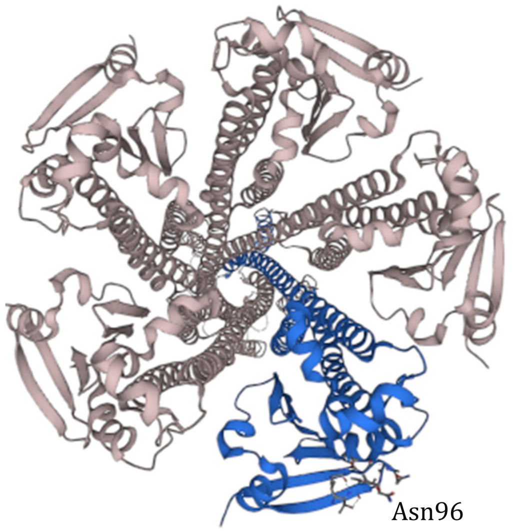


**Supplemental Figure S2B**. Alphafold homology model of MRS2 https://www.ebi.ac.uk/pdbe/pdbe-kb/proteins/Q9HD23/structures , colors indicate confidence of structural assignment of residues from highest confidence (dark blue) to lowest (orange). The two transmembrane helices are at the left and the α/β matrix domain at the right. The proposed *N*-glycosylation site, N96, is highlighted as a ball and stick model, where N96 is part of a β-turn with the side chain amide exposed to solvent. The features highlighted here are present in the recently released cryo-EM structure of human MRS2, pdb8IP3, **https://doi.org/10.2210/pdb8ip3/pdb**. Li, M., et al., *Molecular basis of Mg(2+) permeation through the human mitochondrial Mrs2 channel.* Nat Commun, 2023. **14**(1): p. 4713. Weblink: https://www.ncbi.nlm.nih.gov/pubmed/37543649.

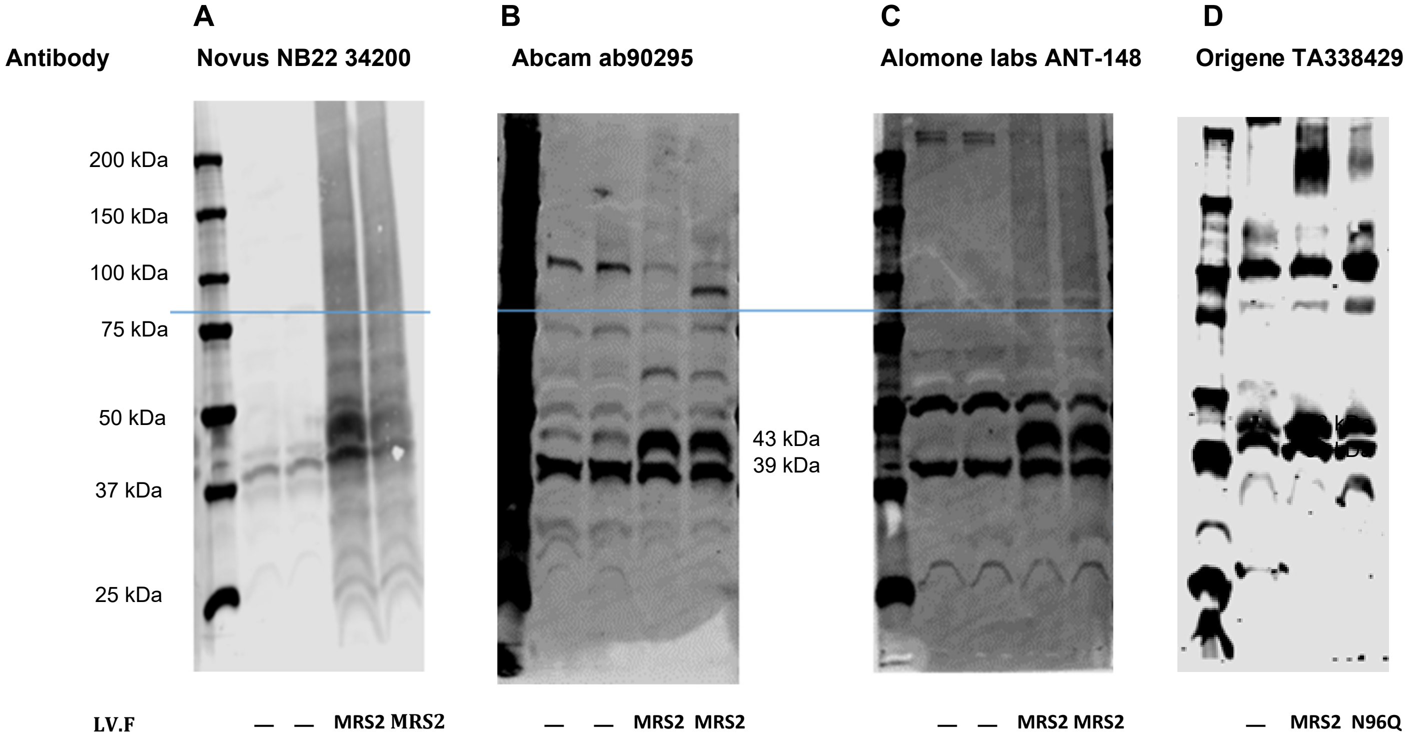


**Supplemental Figure S3**: **Validation of anti-MRS2 antibodies**. Four different antibodies were examined for their sensitivity and selectivity for MRS2: (A) Novus NBP2 34200 (B) Abcam ab90295 (C) Alomone labs ANT-148 (D) Origene: TA338429. In each gel the lane(s) adjacent to the Molecular Weight (MW) markers were mitochondria isolated from mock injected HEK293 cells. In each gel the right two lanes were injected with lentivirus (LV.F) harboring the sequence for WT MRS2 and in (D) lane 3, the MRS2 sequence contained the N96Q mutant. See also for Sigma HPA017642 https://www.proteinatlas.org/ENSG00000124532-MRS2/summary/antibody

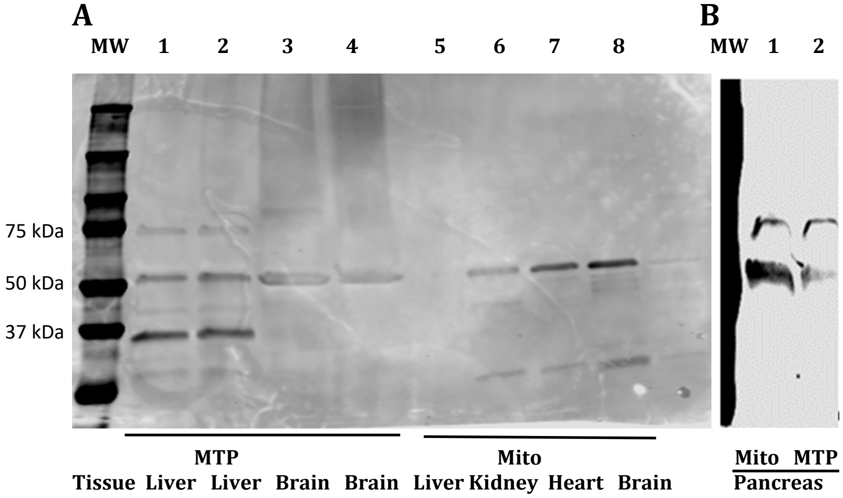


**Supplemental Figure S4.** Anti-MRS2 Immunoblot of mitoplasts (MTP) and mitochondria (Mito) isolated from mouse liver (lanes A1, A2, A5), brain (lanes (A3, A4, A8), kidney (lane A6) and pancreas (B1, B2) uniformly show the presence of an isoform at 52 kDa.
